# Tendon-associated gene expression precedes osteogenesis in mid-palatal suture establishment

**DOI:** 10.1101/2024.05.11.590129

**Authors:** Daniela M. Roth, Jeremie Oliver Piña, Resmi Raju, James Iben, Fabio R. Faucz, Elena Makareeva, Sergey Leikin, Daniel Graf, Rena N. D’Souza

## Abstract

Orthodontic maxillary expansion relies on intrinsic mid-palatal suture mechanobiology to induce guided osteogenesis, yet establishment of the mid-palatal suture within the continuous secondary palate and causes of maxillary insufficiency remain poorly understood. In contrast, advances in cranial suture research hold promise to improve surgical repair of prematurely fused cranial sutures in craniosynostosis to potentially restore the obliterated signaling environment and ensure continual success of the intervention. We hypothesized that mid-palatal suture establishment is governed by shared principles with calvarial sutures and involves functional linkage between expanding primary ossification centres with the midline mesenchyme. We characterized establishment of the mid-palatal suture from late embryonic to early postnatal timepoints. Suture establishment was visualized using histological techniques and multimodal transcriptomics. We identified that mid-palatal suture formation depends on a spatiotemporally controlled signalling milieu in which tendon-associated genes play a significant role. We mapped relationships between extracellular matrix-encoding gene expression, tenocyte markers, and novel suture patency candidate genes. We identified similar expression patterns in FaceBase-deposited scRNA-seq datasets from cranial sutures. These findings demonstrate shared biological principles for suture establishment, providing further avenues for future development and understanding of maxillofacial interventions.

## Introduction

Despite the many advances in understanding of the molecular basis of palatal development, as well as candidate genes contributing to the pathogenesis of orofacial clefting, clinical solutions remain limited and variable in outcome. For example, maxillary skeletal deficiency remains a potential outcome following surgical cleft palate correction. Successful management of transverse maxillary deficiency is dependent upon the intrinsic regenerative capacity of the mid-palatal suture, a fibrous joint connecting the palatine bones. Most commonly, force is applied to the maxilla through orthodontic appliances to expand the palate through techniques such as rapid maxillary expansion (RME), miniscrew-assisted rapid palatal expansion (MARPE), or surgically assisted maxillary expansion (SARME) if the suture is highly interdigitated (Kaya et al., 2023). These tools for orthodontic manipulation of the mid-palatal suture are rapidly evolving, but our understanding of the underlying biology is still incomplete. Cranial sutures are comparatively well-studied with respect to osteoblast and mesenchymal stromal cell behavior under different conditions. The mechanisms enabling their developmental establishment are still under debate. Based on broad biomechanical and osteogenic properties, the mid-palatal suture bears significant resemblance to cranial sutures.

The development of the secondary palate is well studied from the perspective of palatal shelf patterning, elevation, elongation, and mesenchyme fusion. This level of molecular understanding remains incomplete beyond the point of epithelial to mesenchymal transition (EMT) wherein the epithelial seam degrades or transitions to mesenchyme. While the former has been pivotal in understanding the etiology, incidence, and prevention of cleft lip and/or palate (CL/P), the latter is required to address long-term maxillary insufficiency in patients having undergone CL/P repair and complications following routine maxillary expansion (Diah et al., 2007; Gurel et al., 2010; Liberton et al., 2020; Williams et al., 2012; Ye et al., 2015). A similar challenge exists in the field of craniosynostosis management, the premature obliteration of cranial sutures by bone (Stanton et al., 2022). Current treatment begins with excision of the fused suture, with ongoing innovation to prevent re-synostosis. The longevity of CL/P and craniosynostosis repair both depend on eventual establishment of a functional, patent suture with intact mechanosensory, osteogenic, and regenerative capacity. Thus, there is an unmet need for robust understanding of the fundamental principles involved in suture establishment for innovation in both calvarial and palatal clinical intervention.

This study aimed to close gaps in our understanding of mid-palatal suture establishment and function in comparison to cranial suture dynamics. Using a variety of genomic analyses including in situ hybridization, single-cell RNA sequencing (scRNA-seq), and highly multiplex in situ mRNA localization of the palate (10X Genomics Visium, Xenium), we characterized the mid-palatal suture according to cranial suture definitions regarding cellular composition and expression of craniosynostosis risk genes. We then identified a core transcriptomic pattern involved in mid-palatal suture establishment composed of stratified expression of tendon-related genes, Wnt signaling, and extracellular matrix modulators. We confirmed cranial suture expression of the same set of genes using publicly available scRNA-seq datasets (Greg Peter Holmes, 2017; Samuels et al., 2020). Our analysis also identified candidates for maintaining suture patency in the midline mesenchyme of early mid-palatal sutures. This study links molecular understanding of cranial suture biology with that of the mid-palatal suture, drawing connections to enable further innovation in craniofacial research and suture-based clinical interventions.

## Results

### The mid-palatal suture models craniofacial suture establishment

The palate is a continuous structure separating the oral and nasal cavities (*Figure 1A and B*). Following fusion of the mesenchymal compartments of the palatal shelves around embryonic day 15 (e15) (Bush and Jiang, 2012; Lan and Jiang, 2022; Piña et al., 2023), the subsequent stages of palate development involve expansion of the palatine process of the maxilla towards the midline. By postnatal day 0 (P0), closely approximated bone fronts flank the region which will give rise to the mid-palatal suture. The mid-palatal suture undergoes additional changes by P3 which are beyond the scope of this paper, including formation of a temporary cartilaginous structure (Li et al., 2016, 2015). Based on spatial analysis of cell heterogeneity at e15.5, the palatal shelf is composed of distinct cell populations, including mesenchyme, osteoprogenitors (*Cd200, Alpl*), epithelium (*Krt14*), and vasculature (*Pecam1*) (*Figure 1C-E*). The presence of midline mesenchyme separating bone and a bilateral ossification front is a common feature between cranial sutures and the mid-palatal suture, warranting further exploration.

**Figure 1.**
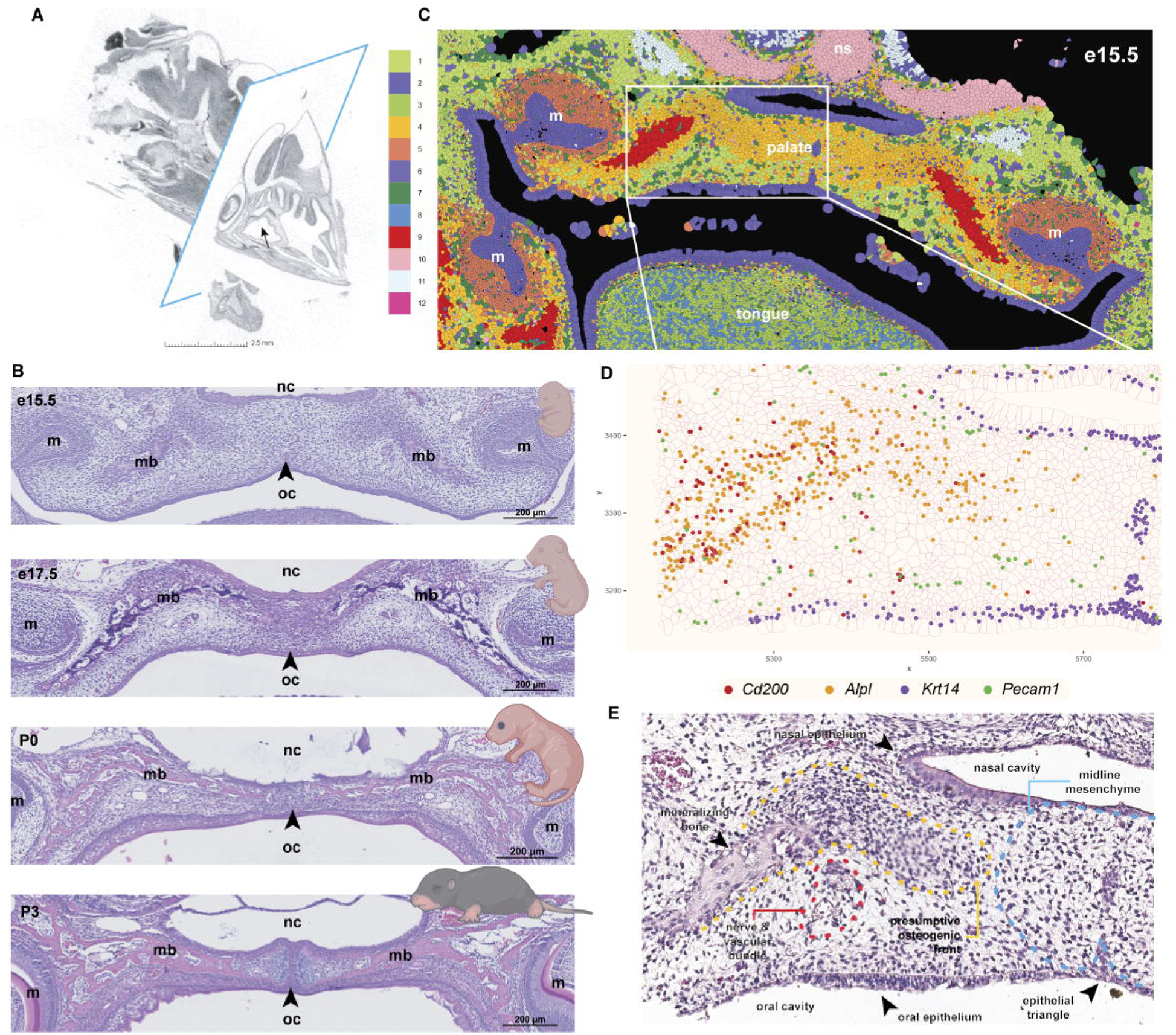
The mid-palatal suture models craniofacial suture establishment. **(A)** Micro-computed tomography reconstruction of iodine-contrasted mouse head at postnatal day 1. 3D image is sliced cross-sectionally in the axial, coronal (blue rectangle), and sagittal planes. Black arrow indicates palatal shelf. Voxel size = 10µm. **(B)** Hematoxylin and eosin-stained 5-10µm thick paraffin sections through the coronal plane of the mid-palatal suture at ages e15.5, e17.5, P0, and P3. Arrow indicates midline mesenchyme. Mouse icons from BioRender. **(C)** 10X Xenium Giotto spatial in situ plot from e15.5 coronal section with cell polygons colored by Leiden clustering algorithm**. (D)** Spatial plot of transcripts for *Cd200* (osteoblasts), *Alpl* (pre-osteoblasts and progenitors), *Krt14* (epithelium), and *Pecam1* (vasculature). Colored circles indicate target genes in legend**. (E)** Annotated H&E of Xenium section in C,D. m: molar; mb: mineralizing palatine process of the maxilla; nc: nasal cavity; ns: nasal septum; oc: oral cavity

### Basic cranial suture composition is consistent with that of the mid-palatal suture

The early (e17.5) mid-palatal suture mesenchyme is flanked by osteogenic fronts, exemplified by a sequential distribution of pre-osteoblasts (*Runx2*), osteoblasts (*Sp7*), and osteocytes (*Dmp1*) similar to cranial sutures (*Figure 2A and B, Supplementary Figure 1*) (Farmer et al., 2021). The mid-palatal suture lies between epithelial layers separating the nasal and oral cavities. In contrast, the frontal suture is only bordered by epithelium ectocranially, with the dura mater in direct contact on its endocranial side. When characterized at the single cell transcriptomic level, cells identified in the e15.5 mid-palatal suture clustered into similar cell type groupings in UMAP space as the e18 frontal suture (*Figure 2C-F*). The Xenium spatial transcriptomic platform was used for spatial contextualization of the identified cell types on e15.5 histological sections of the palate (*Figure 2D*). The top 3 genes identified in each cluster of the e15.5 Xenium mid-palatal suture dataset were also observed in comparable cell clusters from the e15.5 mid-palatal suture and e18 frontal suture scRNA-seq datasets, with similar marker genes for frontal suture mesenchyme as in palate mesenchymal clusters in both scRNA-seq and spatial transcriptomic data (*Figure 2G-I*). For example, *Itm2a*, *Col1a1*, and *Col12a1* were mapped to similar mesenchymal clusters in all three datasets. This established feasibility of cross-analysis of cluster components between the mid-palatal suture and cranial suture for further interpretation.

**Figure 2.**
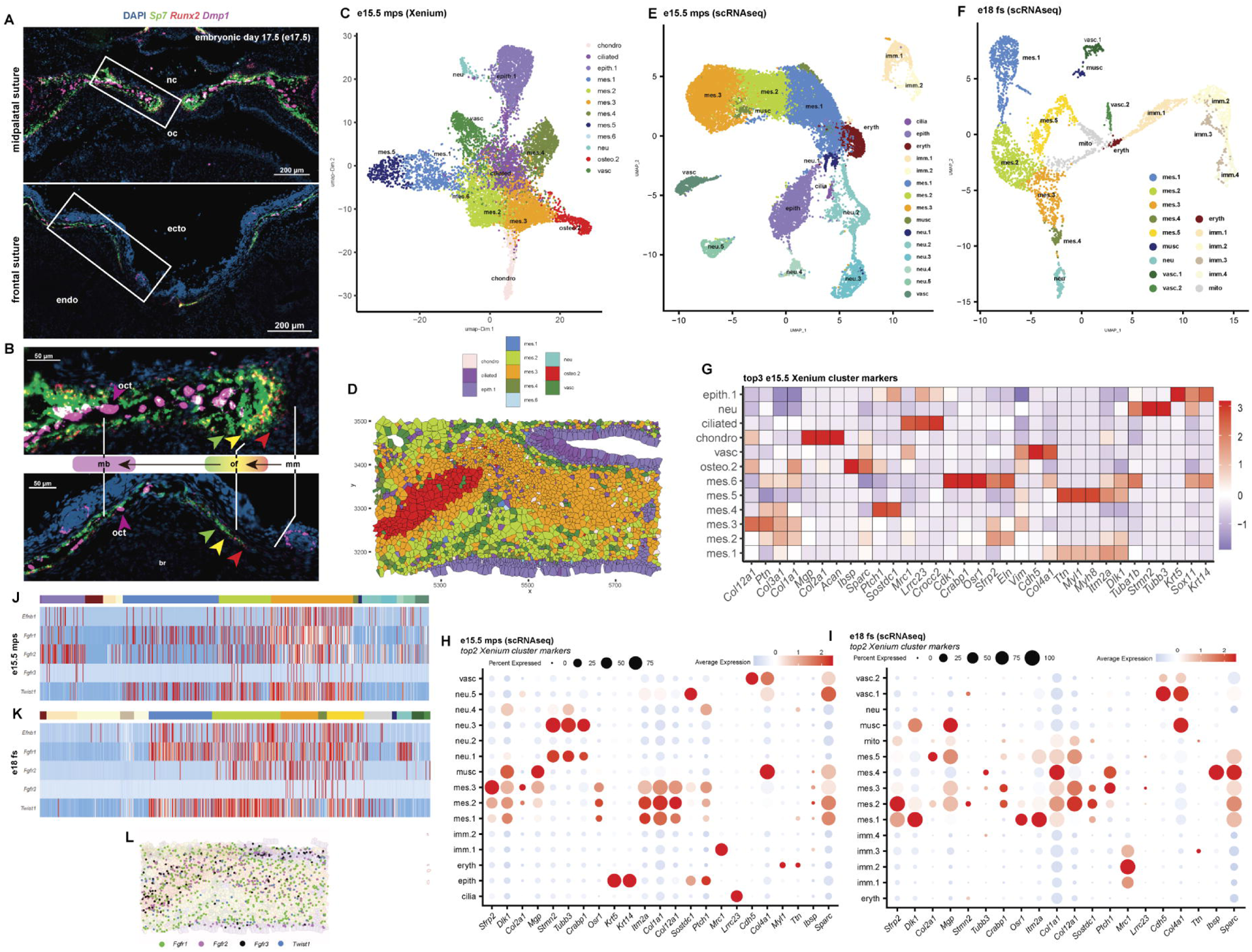
Basic cranial suture composition is consistent with that of the mid-palatal suture. **(A)** RNAscope mRNA detection of osteogenic markers in the mid-palatal suture and frontal suture. **(B)** Magnified region from white box in A, highlighting the gradient of osteogenic differentiation from mesenchyme to bone in both structures. Red arrows indicate *Runx2+* cells, green arrows indicate *Sp7*+ cells, and yellow arrows indicate double positive *Runx2+/Sp7+* osteoblasts. Magenta arrows highlight *Dmp1+* osteocytes. nc: nasal cavity; oc: oral cavity; ecto: ectocranial; endo: endocranial; oct: osteocyte; br: brain. **(C)** Xenium spatial transcriptomic UMAP of e15.5 palate using Giotto analysis pipeline. Leiden clusters annotated based on top gene markers and spatial localization. chondro: chondrogenic; epith: epithelial; mes: mesenchymal; neu: neural; osteo: osteogenic; vasc: vascular. **(D)** Giotto spatial projection of left e15.5 palatal shelf, with polygons labeled according to annotated Leiden clusters (as in C). **(E)** e15.5 palate (mps) single cell RNA sequencing (scRNA-seq) UMAP projection. Seurat clusters annotated based on top genes identified with the FindMarkers function. epith: epithelial; eryth: erythrocytes; imm: immune; mes: mesenchymal; musc: muscle; neu: neural; vasc: vascular. **(F)** E16 frontal suture (fs) scRNA-seq UMAP projection. Dataset downloaded from FaceBase courtesy of Holmes et al. Seurat clusters annotated based on top genes identified as in E. **(G)** Heatmap of top 3 gene markers in each e15.5 Xenium Leiden cluster. mes: mesenchymal; musc: muscle; neu: neural; vasc: vascular; eryth: erythrocytes; imm: immune; mito: mitochondrial. **(H)** Heatmap of expression of top 2 marker genes from e15.5 Xenium clusters in e15.5 mps scRNA-seq dataset. **(I)** Heatmap of top 2 marker genes as in H, according to their expression in e18 fs scRNA-seq dataset. **(J,K)** Heatmaps of expression of genes frequently associated with craniosynostosis in e15.5 mps scRNA-seq clusters (J) and e18 fs scRNA-seq clusters (K). Highest expression in both datasets for all genes appears to be focused around mesenchymal clusters, with the addition of epithelial expression in the e15.5 mps scRNA-seq dataset. **(L)** Spatial expression Giotto plot (spatplot) of four craniosynostosis-associated genes on the left e15.5 palatal shelf, with *Fgfr1* in green, *Fgfr2* in purple, *Fgfr3* in black, and *Twist1* in blue. Polygons are colored according to Leiden clustering. Each point is a detected transcript for the coded gene.

Genes commonly associated with craniosynostosis (premature fusion of the cranial sutures) such as *Efnb1*, *Fgfr1*, *Fgfr2*, and *Twist1* were found in mesenchymal clusters in the mid-palatal suture e15.5 scRNA-seq dataset, with highest expression in mesenchymal cluster 3 (mes.3) (*Figure 2J*). A similar mesenchymal expression is observed in the e18 frontal suture scRNA-seq dataset (*Figure 2K*). *Fgfr3* was infrequently expressed in either dataset, though followed the same general mesenchymal association. Mapping craniosynostosis-associated genes back to the e15.5 Xenium mid-palatal shelf revealed that *Fgfr1*, *Fgfr2,* and *Twist1* were similarly expressed along the expanding osteogenic front (“mes. 3” cluster) (*Figure 2L, Supplementary Figure 2*). In addition, the *Fgfr* expression was also observed in the ciliated nasal epithelium (“cilia” cluster), absent from the frontal suture dataset.

### The palate osteogenic front is preceded by a gradient of spatially correlated genes

With the knowledge that, by birth, the palate midline would closely resemble a patent, approximated suture, we sought to identify transitional gradients or guiding forces already present in the e15.5 palate mesenchyme. To this end, we used bioinformatic techniques to interpret complex relationships between clusters, their localization, and gene expression. First, we generated a Delaunay-triangulated network of correlation for the e15.5 palatal shelf *(Figure 3A)*. Using the Delaunay network as a reference for spatial correlation and the Giotto package’s cell proximity enrichment algorithm, we generated scores for cell proximity and interactions between annotated Seurat clusters, plotted in *Figure 3B*. Focusing on interactions between different mesenchymal clusters, we identified that three of the highest inter-group interactions were between mes.2—mes.4, mes.2—mes.3, and mes.3—mes.4, indicating spatial relationships between the different transcriptionally distinct mesenchymal clusters (*Figure 3C*). We then identified six spatially correlated “meta-feats” using the binSpect rank method to draw connections beyond individual gene cluster to cluster comparison *(Figure 3D)*. This method uses lists of genes to computationally identify correlated patterns of expression and overall trends. Each of the identified meta-feat clusters mapped back to the original tissue in unique patterns, though some general categories were evident such as epithelial localization of patterns 1 and 3 *(Figure 3E)*. Pattern 5 identified a mesenchymal gradient away from pattern 4, with maximal meta-feat expression in the midline mesenchyme. This pattern representing a group of spatially correlated meta-feats connecting bone with midline was particularly interesting as it might offer clues to explain how the advancing osteogenic front and “patent” suture midline mesenchyme interact, both of which are central to successful suture establishment.

**Figure 3.**
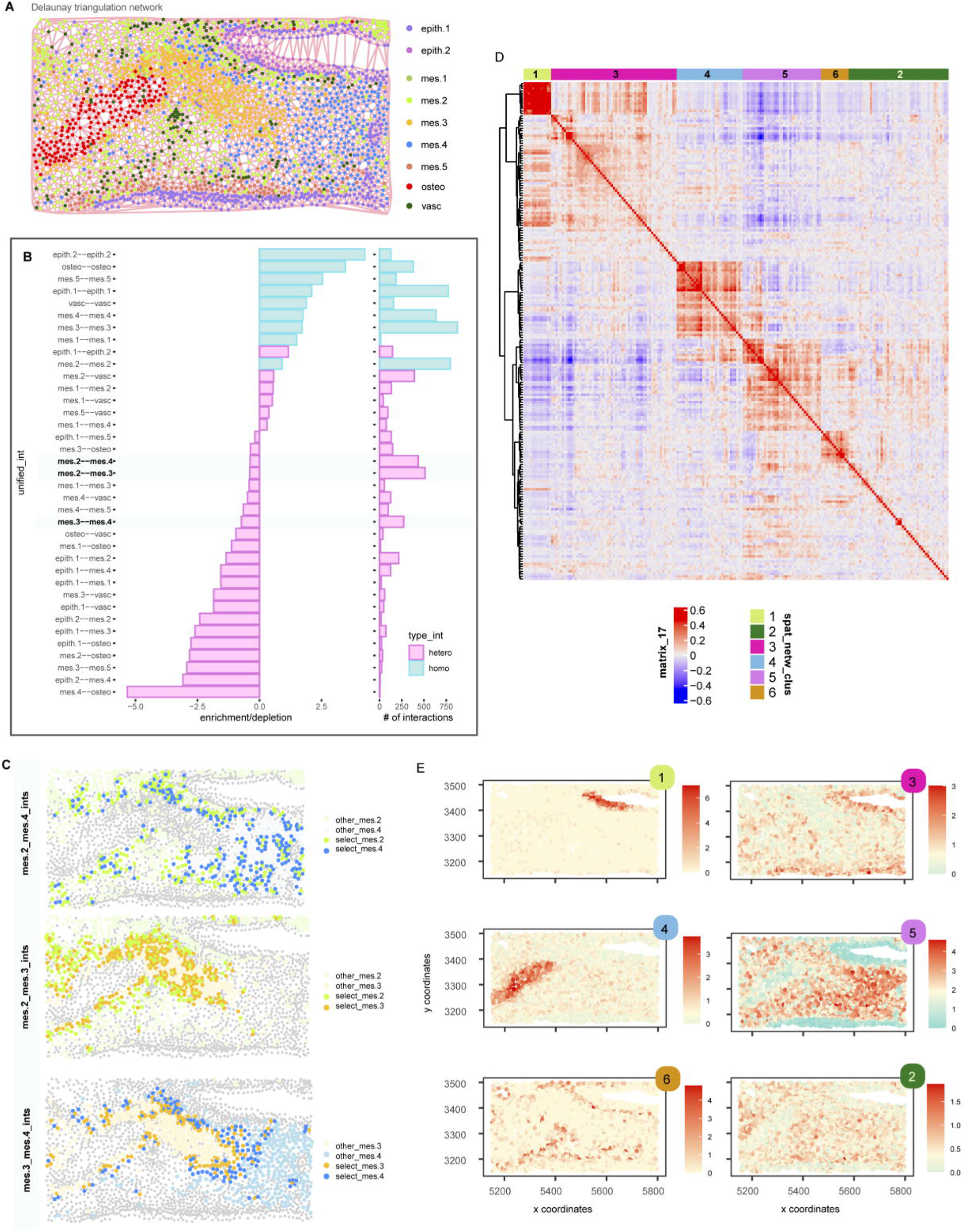
The midline-approaching osteogenic front in the palate is preceded by mesenchyme with a gradient of spatially correlated genes. **(A)** Delaunay network of correlation between clusters in e15.5 left palatal shelf Xenium experiment. Pink lines indicate edges between nodes of the undirected graph output by Delaunay triangulation. **(B)** Cell Proximity Barplot of enrichment or depletion and number of interactions between different annotated clusters at e15.5 (e.g. mes.2—mes.3 indicates spatially correlated interaction between cluster mes.2 and cluster mes.3). Bars are colored according to intra-cluster (homo) or intercluster (hetero) interaction type. Bolded interactions are featured in panel (C). **(C)** Visualization of interacting cells in mes.2—mes.4, mes.2—mes.3, and mes.3—mes.4 interactions. **(D)** Heatmap of clusters generated from spatial co-expression analysis using binSpect rank method. Six spatially correlated patterns of expression were identified. **(E)** Spatial expression of six “metafeats” identified in panel (D).

### Tendon-associated genes are expressed between the midline mesenchyme and palate osteogenic center

At e15.5, a band of stromal cells bridging the mineralizing bone (*Dmp1+*) with the palate midline and its epithelial remnants (*Krt14+*) expressed tendon-associated genes, like *Mkx* and *Chodl (Figure 4A)*. Other tendon- or ligament-related genes identified in cranial suture mesenchyme (Farmer et al., 2021; Holmes et al., 2020) were also enriched in Pattern 5, including *Col3a1, Lum, Six2,* and *Mkx (Figure 4B, Supplementary Figure 3*). *In situ* hybridization of *Sox5* demarcated a heterogeneous population of mesenchyme that was spatially restricted and contained partially overlapping subsets of cells expressing *Col6a2, Col12a1*, and *Chodl (Figure 4C-E). Col6a2* was expressed in the *Sox5+* domain most proximally to the mineralizing bone *(Figure 4D, Supplementary Figure 4*). *Col12a1* was co-expressed by *Col6a2+* cells and extended toward the medial mesenchyme but was notably absent in the most central mesenchyme. *Chodl* was expressed by cells in the superior *Sox5/Col12a1* double positive region surrounding the distinct *Col12a1-* midline mesenchyme.

**Figure 4.**
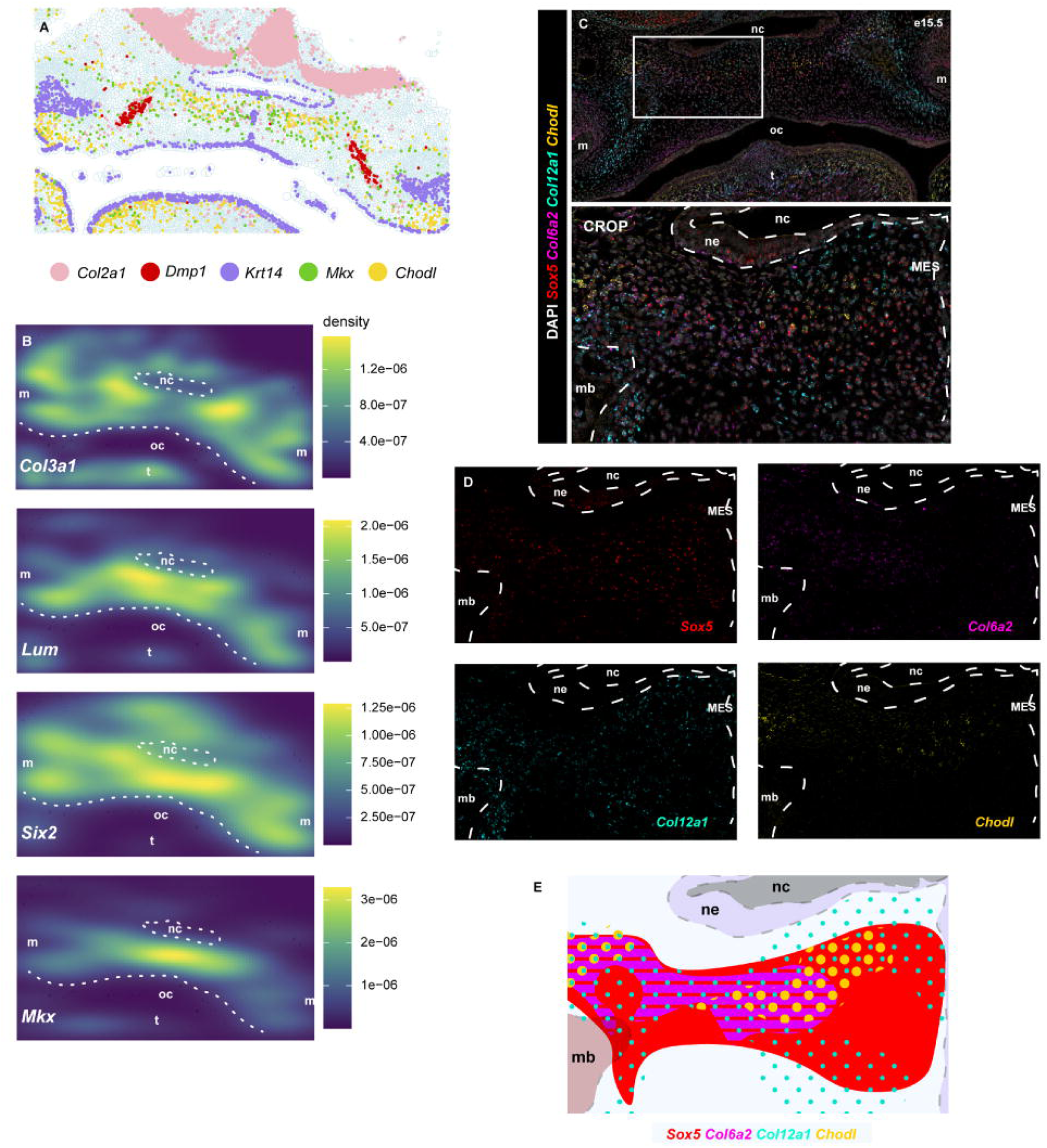
Establishment of the mid-palatal suture involves gradiated expression of tenocyte- and extracellular matrix-associated genes between the midline mesenchyme and osteogenic center. **(A)** Giotto spatial transcript plot contextualizing *Mkx* and *Chodl* in the e15.5 palate, with *Col2a1* mainly marking chondrocytes, *Dmp1* marking the mineralizing palatine process of the maxilla, and *Krt14* marking epithelium. **(B)** Giotto spatial density plots of known tenocyte markers in Xenium dataset: *Col3a1*, *Lum, Six2, Mkx.* Scale indicates density of expression, with dark blue denoting zero and yellow as maximal density for each gene. **(C)** Hiplex RNAscope *in situ* hybridization of probes targeting *Sox5* (red), *Col6a2* (magenta), *Col12a1* (cyan), and *Chodl* (yellow) in the coronal e15.5 palate. DAPI nuclear counterstain is grey. Top panel: palatal shelf overview; white rectangle indicates region of cropped view in lower panel. **(D)** Isolated split channel images for cropped left palatal shelf in C. **(E)** Graphic representation of expression domains of *Sox5, Col6a2, Col12a1,* and *Chodl*, drawn using Adobe Illustrator. oc: oral cavity; m: molar; mb: mineralizing bone; MES: midline epithelial seam ;mc: Meckel’s cartilage; mn: mandible; nc: nasal cavity; ne: nasal epithelium; ns: nasal septum; t: tongue

### Midline mesenchyme marker genes become more restricted in expression over developmental time in the mid-palatal suture and cranial sutures

The e15.5 midline mesenchyme is histologically identifiable as condensed mesenchyme surrounding the formerly continuous midline epithelial seam (MES) *(Figure 5A).* Even at low resolution, the midline mesenchyme clusters into distinct transcriptional identities when compared with other palatal populations (cluster 4) with Visium whole-transcriptome analysis *(Figure 5B).* Genes which were spatially enriched in the palate midline *(Supplementary Files 1-2)*, such as *Sfrp2* and *Tnn*, were also co-expressed in frontal suture scRNA-seq mesenchymal clusters from e16 to P28 *(Figure 5C,D, Supplementary Figure 5*). Wnt signaling has been associated with regulation of tissue differentiation in response to biomechanical stress (Byun et al., 2014; Robinson et al., 2006; Wu et al., 2020). To query Wnt signaling in the context of biomechanically active suture development, we localized Wnt receptor-encoding *Lrp6* and Wnt modulator/osteocyte marker *Dkk1* in the developing craniofacial complex at e17.5 and P3 *(Figure 5E)*. In coronal planes of sectioning, we observed midline mesenchyme expression of *Lrp6* in all visible sutures, including the mid-palatal suture. From e17.5 to P3, expression grew more restricted to a thin, central region of the suture mesenchyme – away from *Dkk1* positive osteogenic cells. Intensity of RNAscope staining suggested that abutting sutures had more focally arranged *Lrp6*+ cells than the overlapping frontal suture.

**Figure 5.**
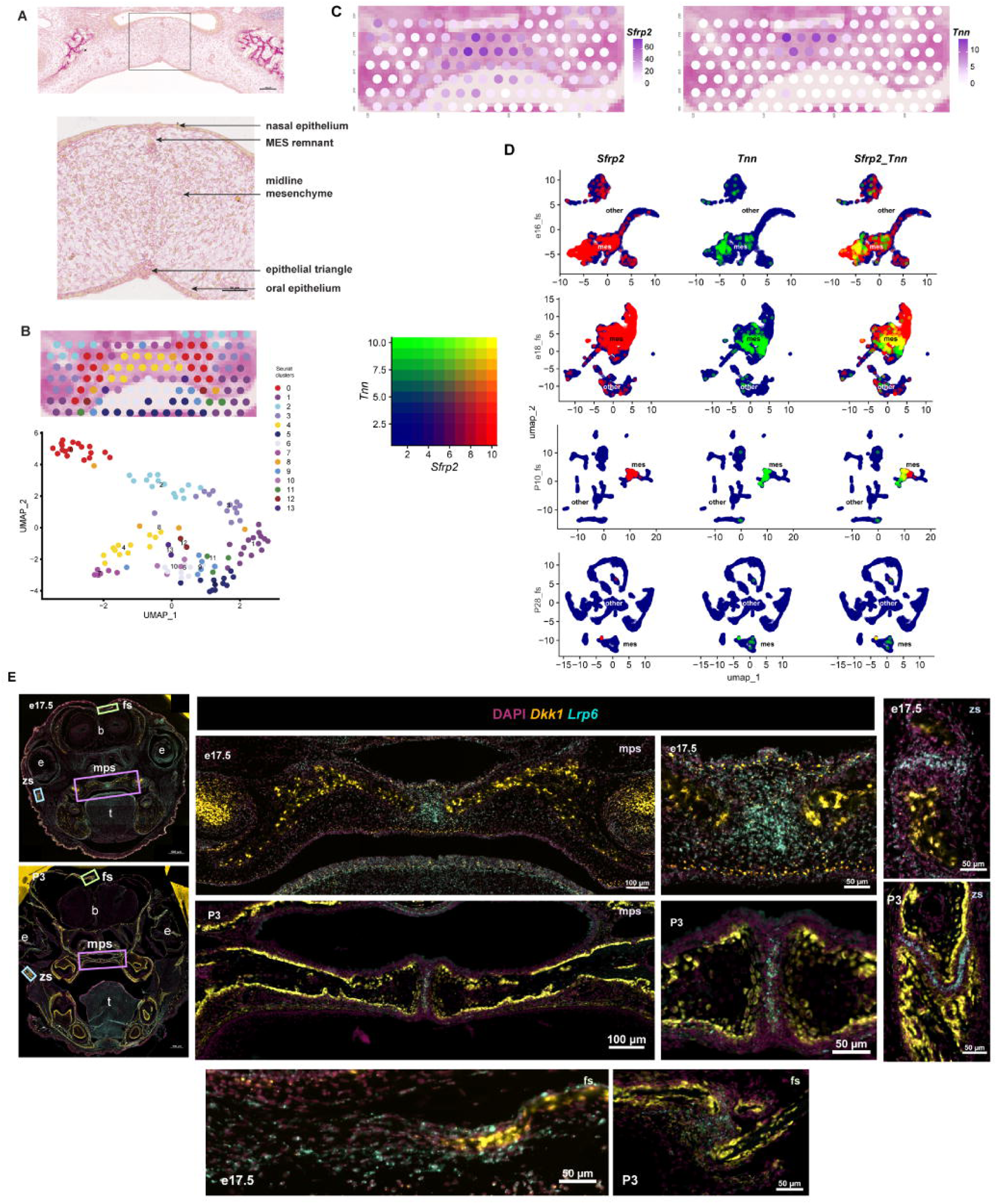
Mid-palatal suture midline mesenchyme marker genes have dynamic expression in cranial sutures, growing more restricted over time. **(A)** e15.5 coronal palate pentachrome histological stain at low magnification, with major components of the midline mesenchyme annotated in lower magnified panel. MES: Midline Epithelial Seam. **(B)** Seurat clusters from graph-based clustering of e15.5 palatal shelf, with yellow (cluster 4) marking the midline mesenchyme on visual assessment. Top panel: spatial dimplot, bottom panel: UMAP. Each circular barcode includes whole transcriptome sequencing information for 3-10 adjacent cells. **(C)** Expression of selected midline mesenchyme markers genes identified from Visium e15.5 cluster 4 targets using the FindMarkers() Seurat function: *Sfrp2* and *Tnn.* **(D)** Joint FeaturePlot expression of *Sfrp2, Tnn,* or both (*Sfrp2_Tnn)* in scRNA-seq UMAP plots generated for e16, e18, P10, and P28 frontal sutures. Red color indicates *Sfrp2* expression, green indicates *Tnn* expression, and yellow points highlight co-expression of both genes. Plots are labeled with categorical annotation of mes (mesenchymal clusters) or other (remaining clusters). Yellow points are mainly found in mesenchymal clusters across all ages. **(E-H)** RNAscope *in situ* hybridization for *Dkk1* (yellow) and *Lrp6* (cyan) mRNA on coronal samples through the middle plane of palate sectioning at e17.5 and P3, overview in E. Note high expression of *Lrp6*, a component of the Wnt signaling pathway, in the midline mesenchyme of all sutures. Colored rectangles indicate regions of magnification for panels F (mid-palatal suture), G (frontal suture), and H (zygomatic suture). DAPI nuclear counterstain is dark violet. b: brain; e: eye; fs: frontal suture; mps: mid-palatal suture; t: tongue; zs: zygomatic suture

### Tenocyte-like cells interface with specific midline mesenchyme

Analysis of the e15.5 midline mesenchyme *in situ* revealed further discrete expression of genes associated with orofacial clefting and Wnt signaling (*Pax9*, for example) *(Figure 6A)*. Interestingly, the TLC extending from the *Dmp1+* bone appeared to directly interface with the *Pax9+* midline. We observed a change in *Col1a1* fiber density and organization at the point of transition from *Col12a1+* TLC to midline mesenchyme *(Figure 6B)*. This boundary was reflected in highly multiplexed *in situ* hybridization between the *Col6a2+* TLC and *Sox9*-enriched midline mesenchyme *(Figure 6C,D)*. We noted *Thbs1* expression along the dense band of collagen fibers. *Scx* was widely expressed by cells throughout the mesenchyme but was highest within the *Sox9+* midline mesenchyme. Stratified expression of extracellular matrix genes radiated from the bone edge (*Mkx, Tnn, Thbs1*) toward midline expression of other important developmental growth factors (*Tgfb2*, *Gsc, Bmp2*) *(Figure 6E,F).* In fact, many core gene regulatory networks of suture development are similarly stratified in expression between the bone and midline, including Bmp, Fgf, Hh, Mmp, Igf, Tgfb, and Wnt pathways *(Supplementary File 3)*.

**Figure 6.**
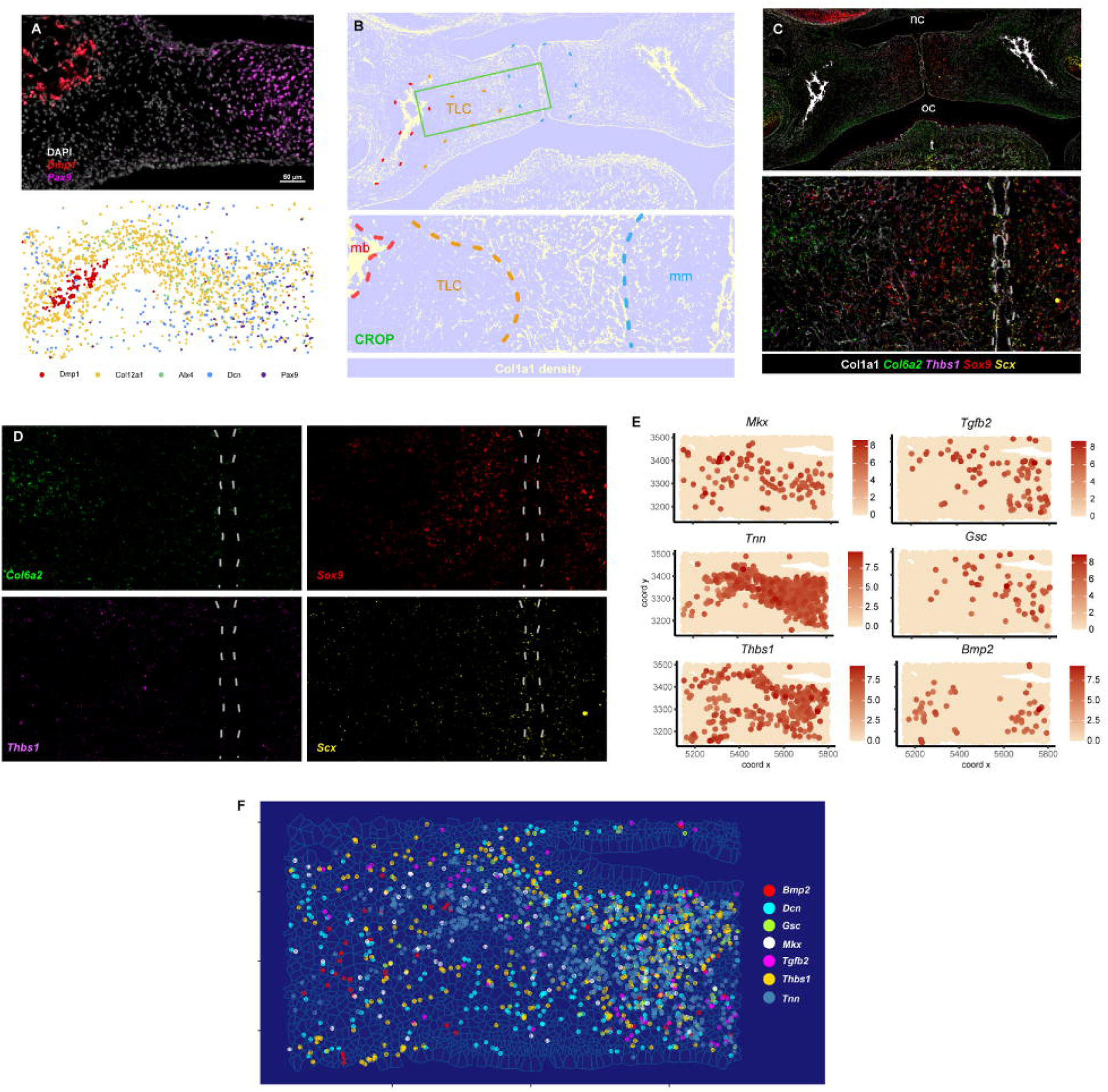
Tenocyte-like cells interface with transcriptionally distinct midline mesenchyme in the palate. **(A)** RNAscope Manual Assay targeting *Dmp1* (representing immature osteocytes and late osteoblasts) and *Pax9*, a transcription factor which is expressed highly in the mid-palatal midline mesenchyme. Lower panel: spatial plot of transcripts identified in e15.5 mid-palatal suture Xenium assay demonstrating gradient of gene expression (*Col12a1, Dcn, Alx4)* between bone (*Dmp1*) and midline mesenchyme (*Pax9*). **(B)** Density visualization of Col1a1 protein immunofluorescent staining in the e15.5 palate, generated using ImageJ. Red dotted outline highlights mineralizing bone (mb), orange denotes the mesial edge of the tenocyte-like population (TLC), and blue outlines the midline mesenchyme (mm). Green rectangle in top panel indicates cropped region below. **(C)** Highly multiplexed *in situ* hybridization of probes targeting *Col6a2*, *Thbs1, Sox9,* and *Scx* assembled with Col1a1 antibody staining on the same FFPE e15.5 coronal section. Bottom panel is cropped to medial edge of left palatal shelf to highlight transitional region bridging *Col6a2* expression with the midline. Grey dotted lines in (C) and (D) indicate the midline epithelial seam. Panels in (D) show individual expression of each gene. **(E)** Giotto spatplot visualization of Xenium targets localized between the mineralizing bone and midline, including TLC, midline mesenchyme, or both (*Mkx, Tnn, Thbs1, Tgfb2, Gsc, Bmp2*). **(F)** Spatial co-localization (Xenium) of *Bmp2, Dcn, Gsc, Mkx, Tgfb2, Thbs1,* and *Tnn* in the context of the e15.5 palatal shelf demonstrating the interface between TLC and the midline mesenchyme. Predicted cell shape polygons are outlined in light blue.

## Discussion

This study assessed the establishment of the mid-palatal suture according to cellular and molecular properties established for the cranial suture, with the aim to identify unique molecular properties of the midline mesenchyme that could mediate suture establishment and patency. Using multimodal transcriptomic techniques, we characterized the architectural and transcriptomic landscape of the palate at e15.5. We identified the presence of cells with a tendon-related molecular signature and provide evidence for a cell hierarchy in the advancing osteogenic front similar to what has been described for cranial sutures. We also characterized the presence of several candidate “patency factors” in the midline mesenchyme of the palate which are similarly expressed in cranial sutures, growing more restricted over time. Our characterization of the shared features of the mid-palatal suture and cranial sutures supports further efforts toward translation between the palate and cranial vault, both clinically and foundationally.

### Tenocyte-like cells form a pseudo-neotendon in the palate

Several patterns of spatially correlated gene expression patterns emerged from our Xenium analysis, but one pattern of “metafeats” in particular seemed to bridge the mineralizing bone with the midline of the palate and showed enriched expression of genes that have been associated with tendons. Tenocytes are specialized tendon fibroblasts which are important for maintenance and transduction of tensile force between bony structures and express *Scx* (Shukunami et al., 2018). It is well established that mesenchymal stromal cell (MSC) differentiation is influenced by mechanical cues. MSCs cultured on a stiffer matrix tend toward myocyte and osteoblast fates (Petzold and Gentleman, 2021), while cyclic stretching can promotes MSC to tenocyte differentiation (Morita et al., 2018). Tenocytes expanded *in vitro*, referred to as “tenocyte-like cells” (TLC), express multipotential genes similar to mesenchymal stromal cells (MSCs) but maintain a potential to differentiate to osteogenic cells (Darrieutort-Laffite et al., 2019; de Mos et al., 2007; Klatte-Schulz et al., 2012). Previous studies have examined the muscular posterior palate at these ages (Grimaldi et al., 2015; Kouskoura et al., 2016; Nara et al., 2017). However, tenocytes have not been characterized in the medial plane of the palate, a region with no direct muscle attachment occurs.

Previous studies using rat embryonic fibroblasts (REF) showed a propensity to condense on soft substrates. This “condensation tendency”, could provide meaningful insight into the formation of the palatal ‘neotendon’ between the stiff palatine process of the maxilla and the soft midline mesenchyme (Xie et al., 2021). In REF, there is an element of tensile prestretch induced by the substrate – an isotropic stretch which causes tissue contraction when released. Based on this, Xie et al conclude that such biomechanical principles are important in defining pattern boundaries, such as between inner and outer boundary cells. A similar principle may underly the patterning of the developing bone front, initiating neotendon formation and creating a stiffer scaffold or path along which osteogenesis may proceed. This possibility is supported by the differential expression of several ECM genes (*Col1a1*, *Col3a1*, *Dcn*, etc). It was also shown that on microfabricated cell sheets cells at the edges of islands along a gradient of traction forces undergo rapid proliferation with observable cell cycle differences, a function which would cause changes in cell signaling (Nelson et al., 2005; Pi et al., 2007). In the e15.5 palate, we observed a collagen-rich neotendon-like cluster of densely packed stromal cells resembling tenocyte-like cells (TLC) based on gene expression with a border of comparatively more sparse cells. This architecture is reminiscent of a neotendon, referred to henceforth as the pseudo-neotendon (or PNT for simplicity). The formation of a PNT could be guided by and/or instructive for the biomechanical nature of the palate, as tendons and ligaments are in other musculoskeletal sites (Liu et al., 2017). This concept underlies recent innovation in regenerative tendon therapy, which may be leveraged for craniofacial applications in mechanically loaded settings (Citro et al., 2023).

### Extracellular matrix in the PNT and cranial suture mesenchyme

A major component of cell migration and force propagation, integrin-mediated activation, is dictated by organization of collagen fibrils (Doyle et al., 2015). The PNT is enriched for genes encoding several collagens. We observed a coordinated pattern of *Col11a1-* and *Col12a1-*expressing cells, as well as proteoglycans and other ECM proteins. Mutations in *Col6a1* and *Col12a1* both affect musculature and result in high arched palate (OMIM 616471, (Baker et al., 2005)). *Col11a1,* most significantly decreased in *Scx-/-* tendons (Liu et al., 2021), is associated with cleft palate in Pierre Robin and Stickler Syndromes (Lavrin et al., 2001) in addition to reports of nonsyndromic CL/P prevalence in COL11A1/2 deficient patients (Guo et al., 2017; Melkoniemi et al., 2003). It should be noted that clefts observed in PRS are usually due to inability of the tongue to contract preventing timely palatal shelves elevation. Interestingly, tongue contraction is dependent on muscle attachment to the mandible, which appears to be compromised in several Pierre Robin Sequence (PRS) mouse models (Kouskoura et al., 2016, 2013; Lavrin et al., 2001). We observed focal expression of genes encoding key proteoglycans that are associated with early palatogenesis within and directly surrounding the PNT, including *Lum* and *Dcn.* Defects in these proteoglycans also have profound effects on tendons with implications in clefting (Hammond et al., 2018; Liu et al., 2021). Previous cranial suture sequencing has shown presence of tendon-associated genes like *Scx, Mkx, Lum, Fmod, Dcn,* and *Col12a1* in the suture mesenchyme at E16.5 and E18.5, suggesting a mechanoresponsive role (Holmes et al., 2020). Farmer et al. discuss and characterize ‘ligament-like mesenchyme’ cell populations in the developing calvarial sutures as being related to suture flexibility, or speculatively playing a role in mechanotransduction with relevance to brain growth (Farmer et al., 2021). These tendon/ligament-like populations were reported to express *Tnmd, Chodl, Scx,* Mkx – all of which were also identified in the TLC herein. Our study supports a central role for tenocyte-like cells in both cranial sutures and the developing mid-palatal suture. Both structures share several features beyond the presence of TLC, such as tensile loading and rapid MSC-related changes allowing coordinated osteogenesis.

### Maintenance of suture patency

An outstanding question is which factors are responsible for maintenance of suture patency. This is relevant, as re-synostosis following suture repair is often observed indicating that re-opening of a suture may not be sufficient to establish all suture characteristics. For a reconstructed suture to be fully functional, it must replicate the *in vivo* signaling environment. Similarly, the mid-palatal suture upon CL/P repair, or even surgically assisted rapid maxillary expansion, should exhibit normal mechanoreception, osteogenic differentiation, and resilience to forces/mechanical disruption. We found a transcriptional identity of midline mesenchyme of the palate characterized by genes associated with extracellular matrix assembly and composition, growth factor signaling, and transcriptional control. *Lrp6* and *Pax9* marked the definitive midline mesenchyme after e15.5. *Lrp6,* which encodes one of two transmembrane Wnt signaling receptors, was similarly expressed in the midline mesenchyme of cranial sutures and showed a notable compression of its expression domain towards the center of the suture mesenchyme in older mice. This finding was interesting to us for two reasons. First, it confirmed that the mid-palatal suture shares key mesenchymal characteristics with other cranial sutures. Also, its continuous yet increasingly restricted expression in the compressed center of the midline of sutures invoked the concept of suture patency. Interestingly, the zygomatic and mid-palatal sutures had higher and more focal expression of *Lrp6* than the frontal sutures – perhaps reflective of the overlapping anatomy of the frontal bones. Furthermore, we identified expression of *Pax9* in a similar expression domain as *Lrp6*. *Pax9* has been associated with CP; however, due to the severe phenotype observed following its deletion, late embryonic to early postnatal studies have not been performed (Ichikawa et al., 2006; Jia et al., 2020; Sweat et al., 2020). Induction of *Pax9* in the dental mesenchyme during mesenchymal condensation has been reported, linking its expression to cellular compression resulting in changes in extracellular matrix composition (Mammoto et al., 2015). We found a spatial relationship of *Pax9* with PNT marker genes in the palate, forming a gradient between the mineralizing palatine process of the maxilla and the *Pax9/Lrp6*-expressing midline mesenchyme. We hypothesize that the induction of midline mesenchyme-enriched factors, like *Lrp6, Tnn,* or *Pax9*, is a result of cell compression due to the approximation of extracellular matrix-enriched osteogenic fronts. The relationship between mesenchymal cell compression, suture establishment, and patency is not yet established (Alves-Afonso et al., 2021) and more detailed study of these factors may be informative.

### Clinical implications

Mid-palatal suture establishment is complex, which must be considered when developing clinical interventions involving this structure. The integration of biomechanical stimuli (mastication, suckling, brain expansion), osteogenesis, odontogenesis, and midfacial growth should be central to these considerations. Cell differentiation is guided by several signaling networks, including WNT, BMP, TGFs, Hh, and FGF pathways. Development and maintenance of the heterogeneous components in the developing palate such as mineralizing bone, vasculature, nerves, epithelium, and cartilage likely not only involves coordinated signaling for lineage differentiation (Chen et al., 2021; Han et al., 2018; Novoseletskaya et al., 2023; Picoli et al., 2024; Tower et al., 2021), but must also include mechanosensitive elements. Thus, it is important to consider how changes in the mechanical environment following surgical intervention might alter the growth potential of the target tissue. Our study brings to light pan-suture involvement of tendon-like cells in suture establishment and identifies as series of novel and shared midline mesenchyme markers. Our data supports a close physical relationship between expanding palatine bone and the patent midline mesenchyme involving a neotendon-like structure, which may be a critical component for formation of a functional suture. It has been noted that even superficial disruption of the mid-palatal suture impedes midface expansion, and midface hypoplasia is still a potential outcome following CL/P repair (Celie et al., 2024; Freng, 1978; Fudalej et al., 2008; Li et al., 2015). The use of MSC-centered/derived therapies for bone regeneration and tendon healing, especially with adjunctive indiscriminate application of rhBmp2, should be revisited under the lens of our findings (Makar et al., 2021; Shen et al., 2022; Takagi & Urist, 1982).

## Materials and methods

### Key resources table

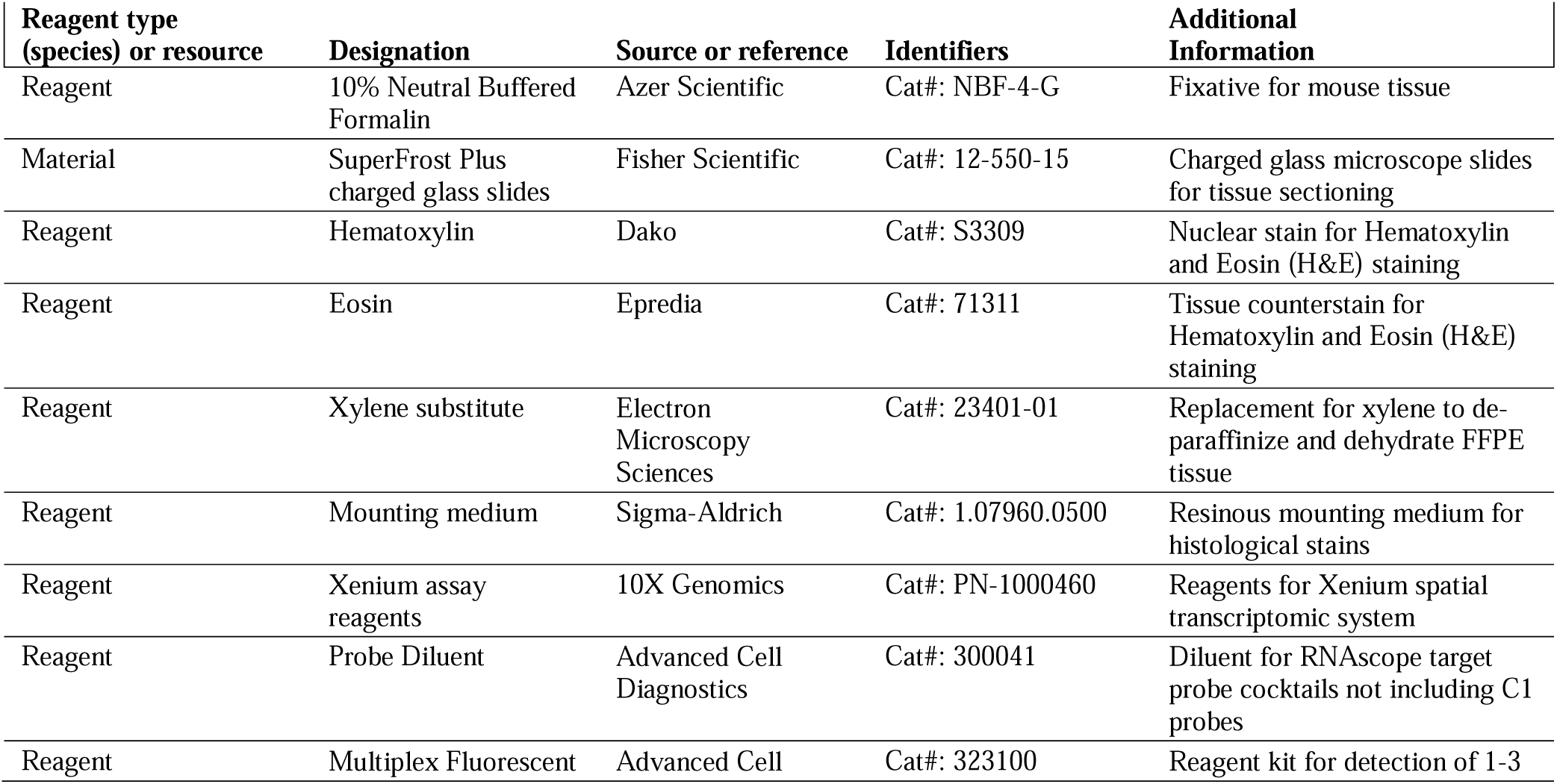
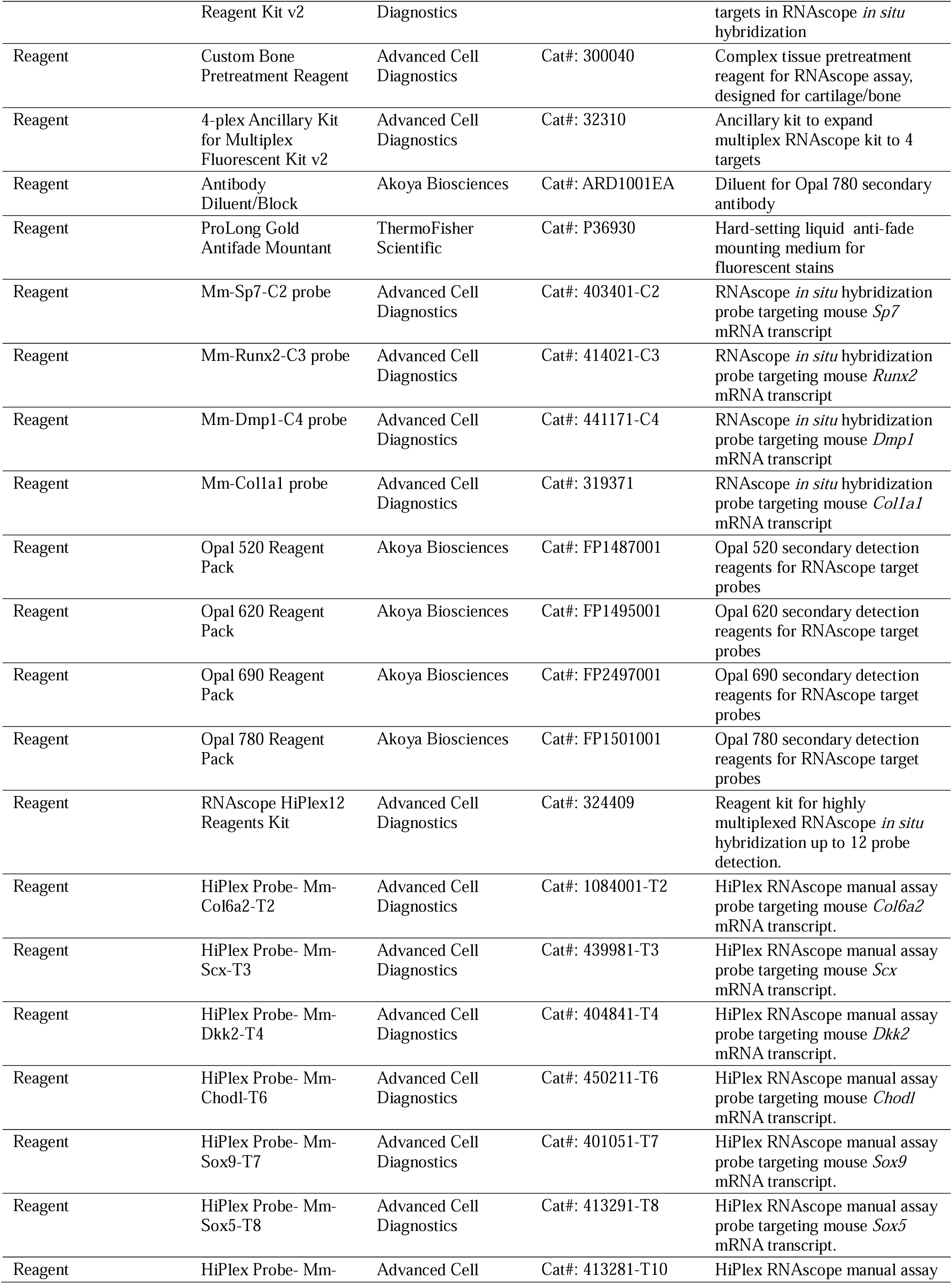
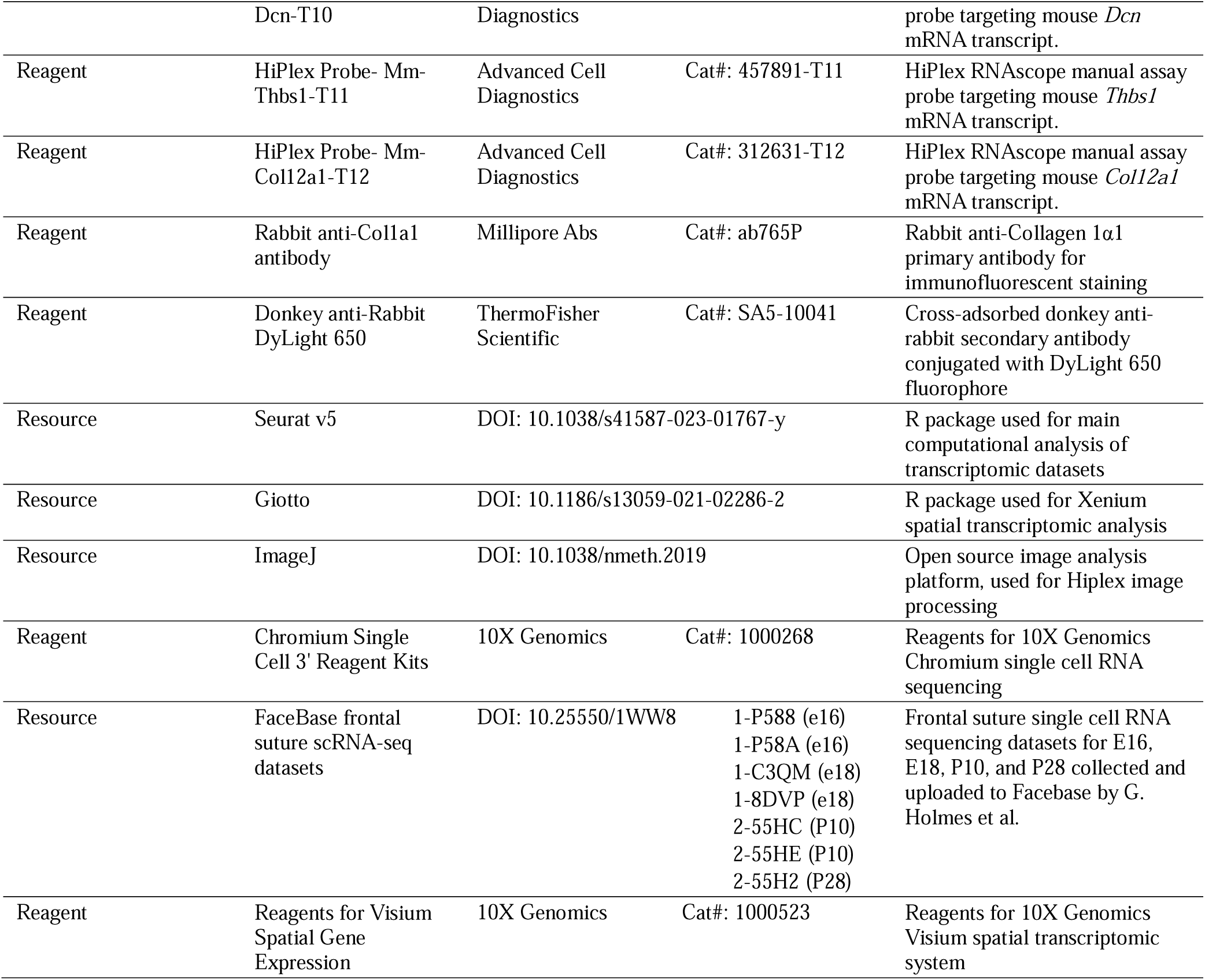

### Animal breeding and husbandry

All mice were of a C57Bl/6 *Mus musculus* genetic background (JAX stock #000664). Mice were housed in controlled conditions with a regular light cycle and free access to water and food at the National Institutes of Health in Bethesda, MD. All animal experiments were approved by the National Institute of Child Health and Human Development Animal Care and Use Committee (ASP #21-031). Timed pregnancies were dated day 0.5 for plug date.

### Tissue processing and embedding

Dams were sacrificed by CO_2_ inhalation and subsequent cervical dislocation. Skinned heads were fixed using 10% neutral buffered formalin at 4°C. Heads were decalcified with 0.5M EDTA, pH 7.4 overnight. After sufficient decalcification, tissue was pre-soaked in 50% ethanol and processed for paraffin embedding in tissue cassettes using a cabinet HistoCore tissue processor according to the following settings: 70% EtOH (1hr), 95% EtOH (1hr), 2X 100% EtOH (1hr each), 3X xylene (1hr each), 2X paraffin (1hr each, with vacuum). Cassettes were then transferred to a 50-65°C paraffin bath for embedding at the Leica paraffin embedding station. Blocks were cooled at 4°C prior to sectioning.

### Histology

Paraffin blocks were sectioned using a semi-automated Leica HistoCore Autocut at 5-10µm thickness onto Superfrost^TM^ Plus charged glass slides (Fisher Scientific, #12-550-15). For all histological stains, slides were deparaffinized and rehydrated as described in (Roth et al., 2022). Briefly, wax was melted at 60°C and slides were washed in a series of xylene and graded EtOH dilutions to water. For hematoxylin and eosin staining, nuclei were stained with hematoxylin then counterstained with eosin, dehydrated, and coverslipped with toluene-based mounting medium. For pentachrome staining, slides were treated with 6% nitric acid, dipped in toluidine blue (0.5g in 100mL ddH_2_O, pH 1-1.5), and incubated in picrosirius red solution with agitation for 10 min or until bone was stained red. Slides were then dehydrated and mounted.

### *In situ* hybridization

ACD Manual RNAscope assays for one- to four-plex *in situ* hybridization were performed as previously described (Piña et al., 2023), using the probes designated in the key resources table above. For Manual Assay RNAscope HiPlex and subsequent immunofluorescent staining, the assay was conducted following the standard protocol provided by the manufacturer, employing the RNAscope HiPlex12 Kit (Advanced Cell Diagnostics Cat. #324409). The gene targets included *Col6a2*, *Scx, Dkk2, Chodl, Sox9, Sox5, Dcn, Thbs1,* and *Col12a1*. Five-micron FFPE tissue sections were baked at 57°C for 30 minutes, followed by deparaffinization in xylene substitute (Electron Microscopy Sciences, Cat. # 23401-01) and dehydration in an ethanol series. Custom pre-treatment reagent treatment was applied for 30 minutes at 40°C, and then hydrogen peroxide (Advanced Cell Diagnostics Cat. # 322335) treatment was performed at room temperature for 10 minutes. Probes for the nine target genes were hybridized for 2 hours at 40°C. Subsequently, the first round of hybridization attached three fluorophores to the first three of the nine target genes (T2-T4). After hybridization, the sections were counterstained with DAPI for 10 minutes and mounted with ProLong Gold Antifade Mountant for imaging using an Eclipse E400 microscope (Nikon, Tokyo, Japan). Following imaging, coverslips were removed using 4X SSC buffer, and fluorophores were cleaved using the provided cleaving solution. A new set of fluorophores targeting the next three genes (T6-T8) were hybridized onto the tissue sections, followed by another round of DAPI counterstaining. Imaging was repeated as previously described. This process was repeated for the remaining genes (T10-T12), capturing images of all nine target genes. T1, T5, and T9 channels were not used for hybridization to allow for autofluorescent calibration in ImageJ using FITC fluorescence.

Immunofluorescence labeling of Col1a1 protein was performed on the same tissue sections following cleavage of fluorophores from the last round of the HiPlex Assay. After removing the coverslips once again, tissue sections were briefly washed in 0.5% Tween-20 in TBS (TBST) and blocked using 10% donkey serum and 1% BSA in TBS-Triton (0.025% Triton X-100) at room temperature for 1 hour with gentle agitation. Slides were then incubated with rabbit anti-Collagen 1α1 primary antibody (Millipore Abs ab765P, 1:100) diluted in blocking buffer at 4°C overnight with gentle agitation. After washing in 0.5% TBST, slides were incubated for 1 hour at room temperature in donkey anti-rabbit DyLight 650 (Thermo Fisher #SA5-10041, 1:500) secondary antibody diluted in blocking buffer. Following three washes in 0.5% TBST, slides were counterstained with DAPI and coverslipped using ProLong Gold Antifade Mountant and slides were imaged. After imaging, coverslips were removed in 4X SSC buffer, tissue sections were briefly washed in 0.5% TBST, and stained with hematoxylin and eosin (H&E) for morphological analysis. Images from each round of RNAscope, immunofluorescence, and H&E staining were processed in ImageJ for background removal, autofluorescence subtraction, alignment and scaling, and assembly of probe combination images.

### Single cell RNA sequencing

For single cell RNA sequencing (scRNA-seq), mid-palatal sutures were dissected with surrounding palate tissue from embryonic day 15.5 (e15.5) or P1 littermates and snap frozen. Tissue handling and RNA sequencing was performed at the NICHD Molecular Genomics Core. Cellranger output files were then used for further downstream computational analysis. In addition to newly generated RNA sequencing data, this project made use of online Facebase scRNA-seq repositories, specifically for frontal sutures from the Holmes group (Greg Peter Holmes, 2017; Samuels et al., 2020). For comparison of mesenchymal and non-mesenchymal clusters in Figure 5, we integrated two bioreplicates each for E16, E18, and P10. All other analysis was done using single replicate datasets.

### Spatial transcriptomics

Freshly cut 5-10µm-thick paraffin sections were used for both 10X Genomics Visium and Xenium spatial transcriptomic assays. Multiple sections were placed within a defined fiducial frame (11mm^2^ for Visium, 12mm by 24mm for Xenium) and baked briefly at 56°C. Slides were processed according to manufacturer-provided protocols, with high resolution brightfield imaging at their conclusion.

### Computational analysis

All computational analysis was done in the R programming environment, mainly using the Seurat v4 and 5 packages (Hao et al., 2024, 2021). Single-cell RNA-sequencing and Visium datasets were primarily processed with Seurat functionality, while Xenium was analyzed according to Giotto vignettes. Other packages used for pre-processing and visualization included PopsicleR (Grandi et al., 2022), Giotto (Dries et al., 2021), and SCpubr (Blanco-Carmona, 2022). Up to date code is located at github.com/dmartaroth. Annotation was done manually based on marker genes from the literature.

## Acknowledgements

The authors would like to thank Parna Chattaraj for her invaluable laboratory technical support and guidance for this project, Dr. Vardit Kram at the NIDCR for CT scanning support, Dr. Jacqueline Tabler for her advice, Dr. Edward Mertz for the use of his microscope, and Dr. Vincent Schram for his expert microscopy at the NICHD microscopy core. This work utilized the computational resources of the NIH HPC Biowulf cluster (http://hpc.nih.gov). DMR and JOP were both supported by National Institute of Child Health and Human Development Pre-Doctoral Intramural Research Training Award (IRTA) Traineeships.

## Additional Information

## Funding

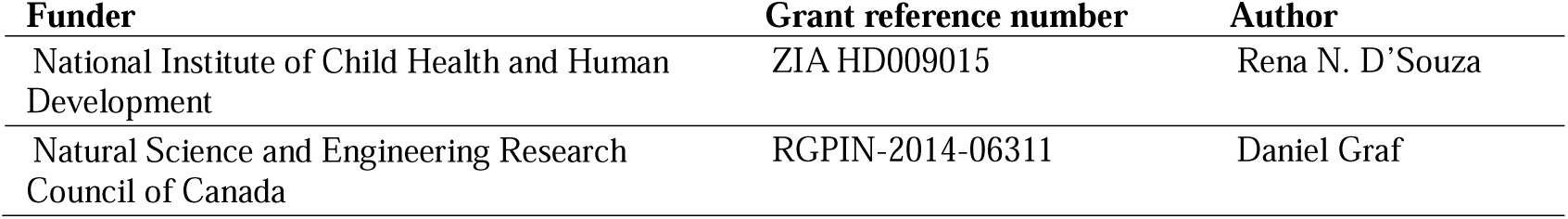

## Author contributions

Daniela Marta Roth, Conceptualization, Methodology, Investigation, Data curation, Writing – original draft, Writing - revisions; Jeremie Oliver Piña, Conceptualization, Investigation, Methodology, Writing - revisions; Resmi Raju, Investigation, Methodology, Writing - revisions; James Iben, Methodology, Data curation; Fabio Rueda Faucz, Investigation, Methodology, Resources; Elena Makareva, Investigation, Methodology; Sergey Leikin, Methodology, Data curation, Resources; Daniel Graf, Supervision, Writing - revisions, Resources, Funding acquisition; Rena N. D’Souza, Conceptualization, Supervision, Resources, Funding acquisition. All authors reviewed the manuscript.

