## Supplementary Figures for "Tendon-associated gene expression precedes osteogenesis in mid-palatal suture establishment"

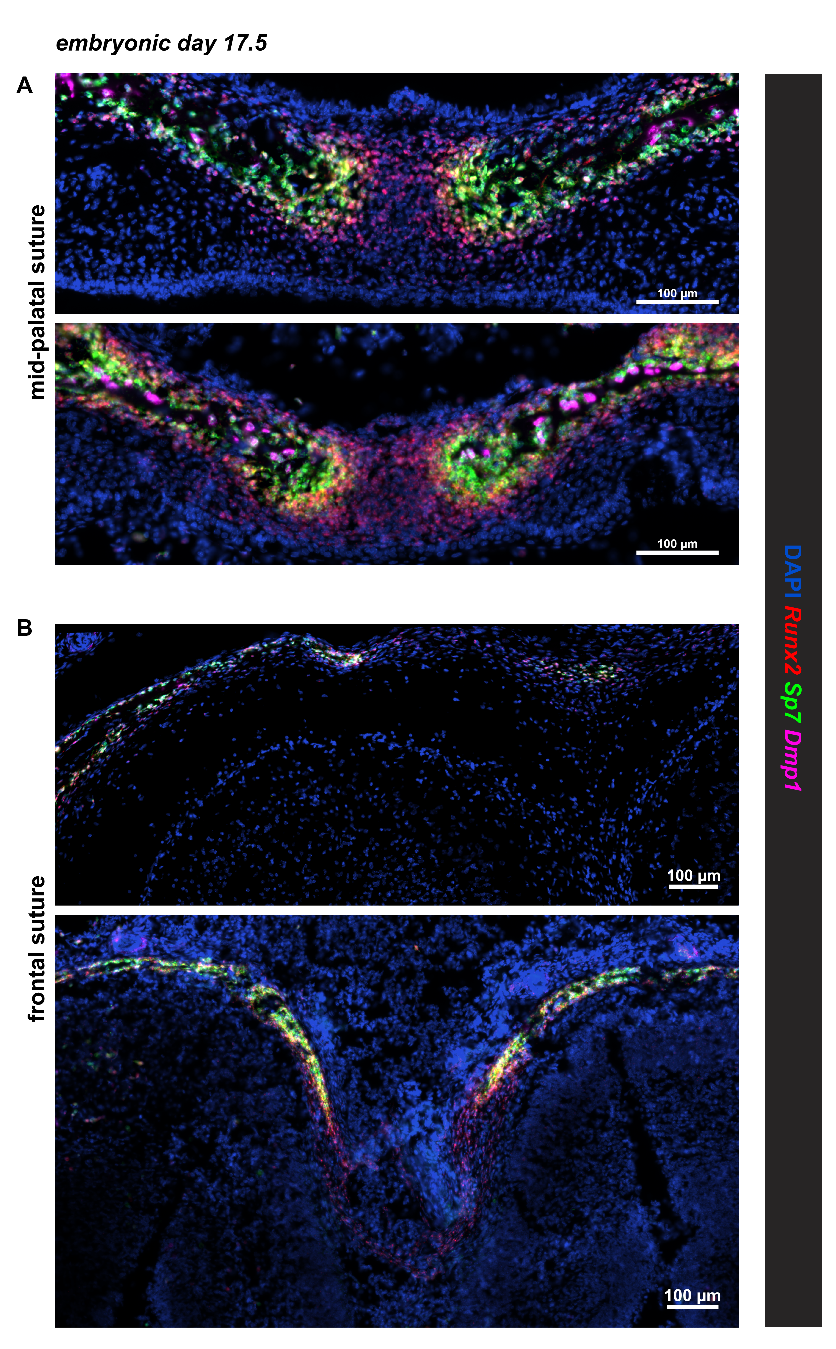

**Supplementary Figure 1. Expression of *Runx2*, *Sp7*, and *Dmp1* in the mid-palatal and frontal sutures.** RNAscope Manual Assay *in situ* hybridization on coronal FFPE embryonic day 17.5 (e17.5) sections in biological duplicate across the mid-palatal suture **(A)** and frontal suture **(B)**.

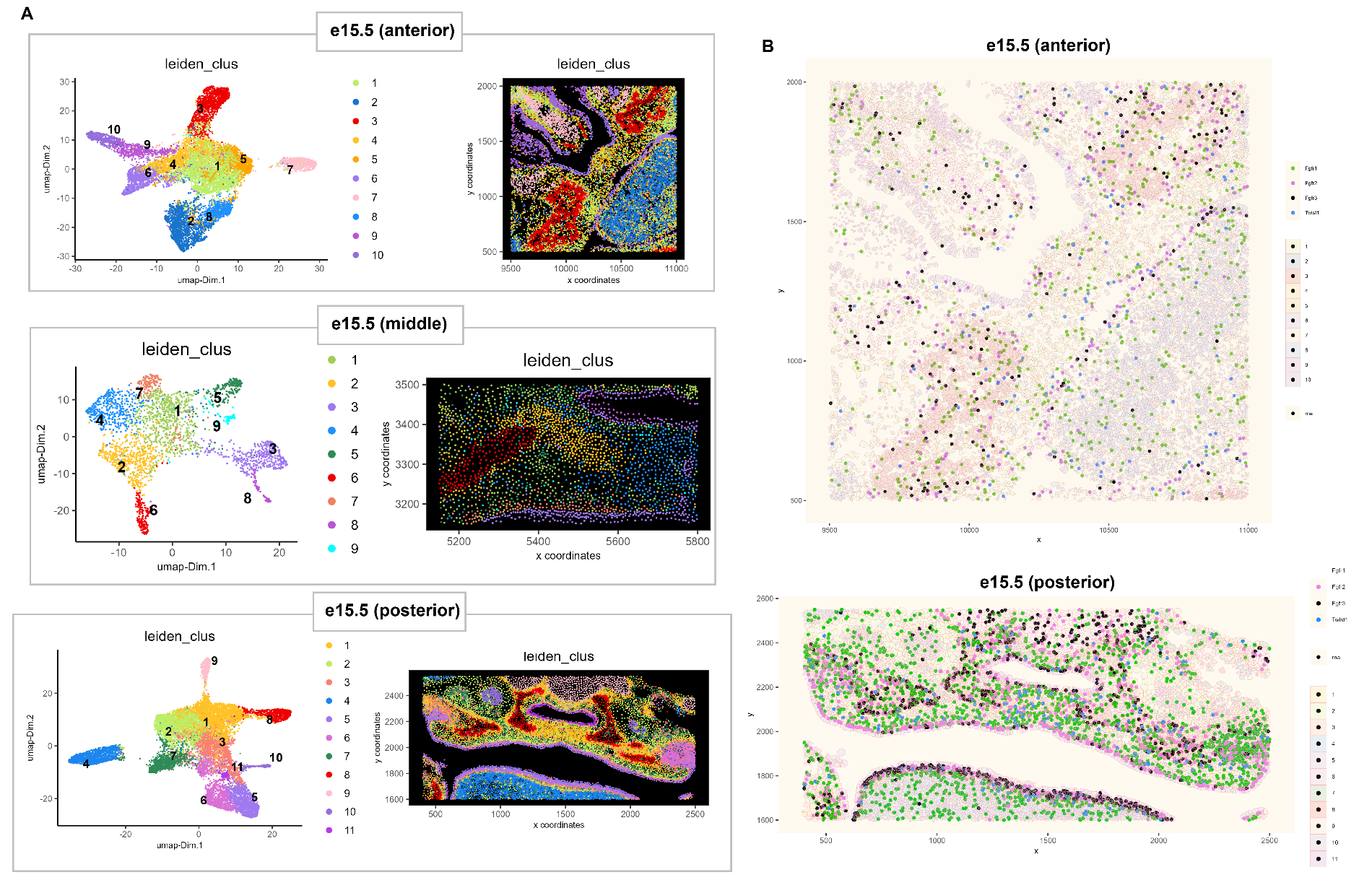

**Supplementary Figure 2.** *Expression of craniosynostosis genes in anterior and posterior planes of the palate.* **(A)** Giotto SpatDimplot comparing leiden clustering results for anterior, middle, and posterior Xenium sections of the embryonic day 15.5 palate in separate mice. **(B)** Expression of craniosynostosis-related genes (*Fgfr1/2/3, Twist1*) in anterior and posterior mid-palatal suture planes.

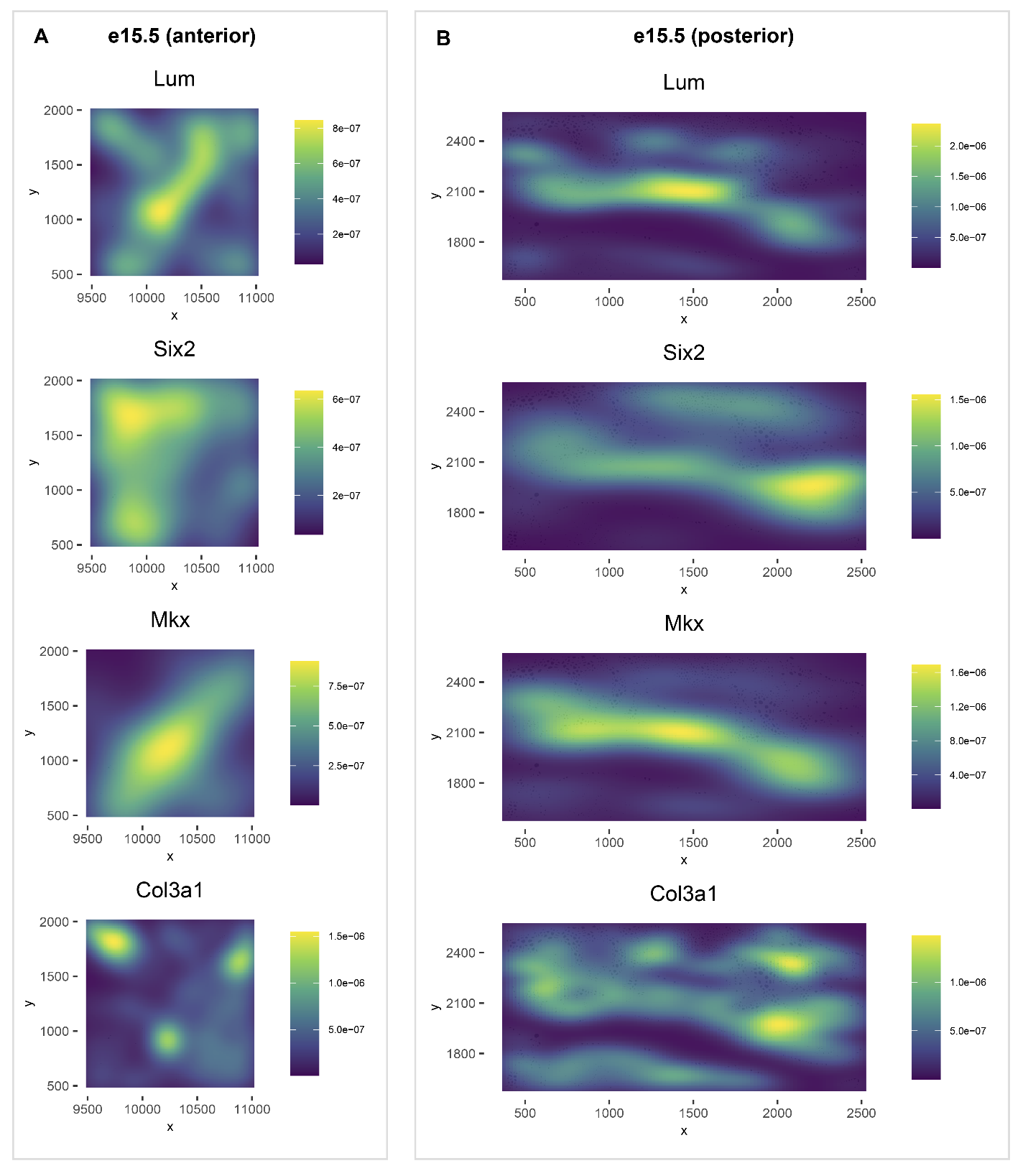

**Supplementary Figure 3. Expression density of tendon-associated genes in anterior and posterior planes of the embryonic day 15.5 palate.** Spatial density plot of *Lum, Six2, Mkx,* and *Col3a1* in anterior **(A)** and posterior **(B)** sections shown in Supplementary Figure 2.

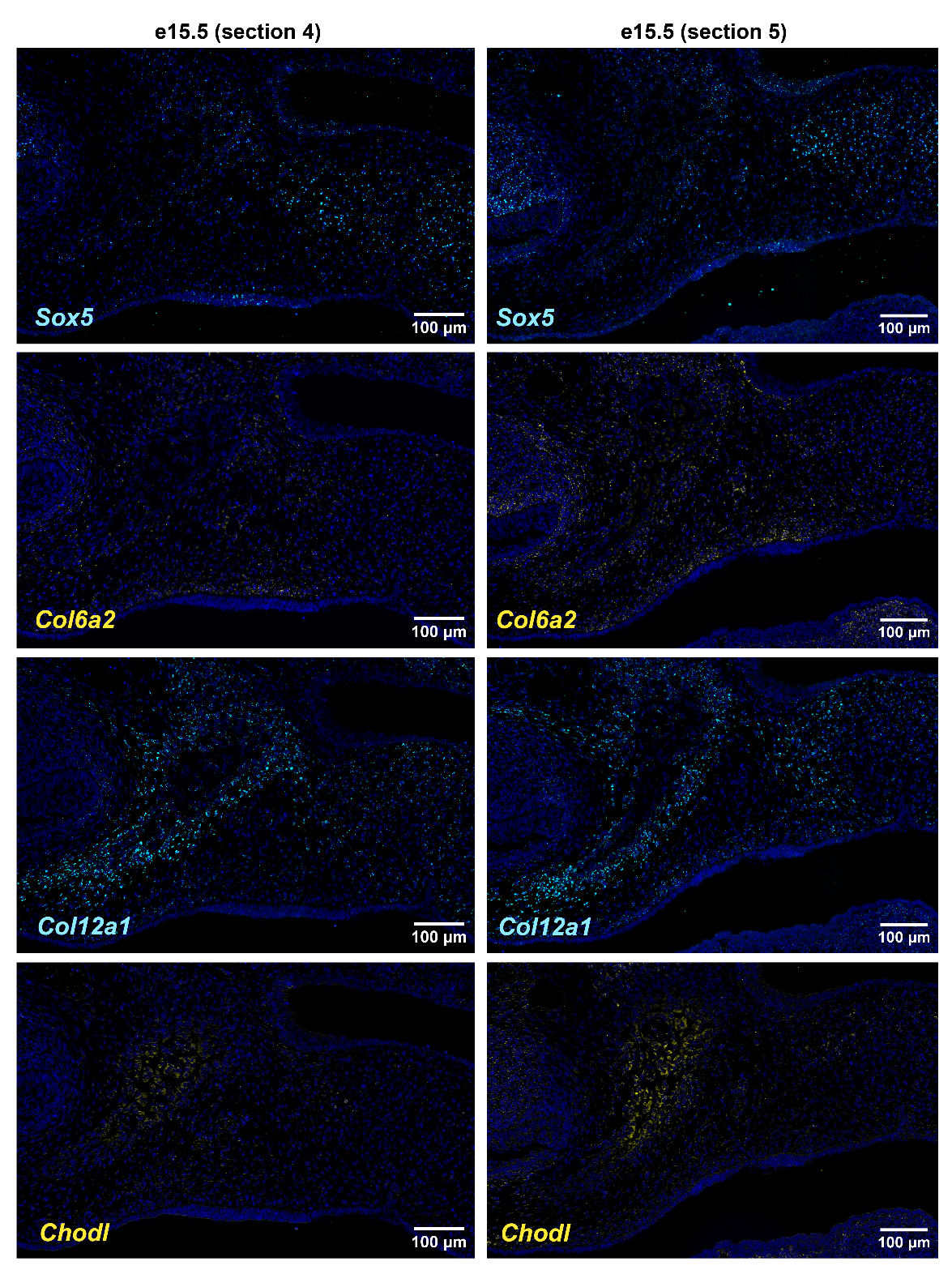

**Supplementary Figure 4. RNAscope Hiplex hybridization of tendon-associated genes in biological replicates.** mRNA localization using highly multiplexed hybridization for *Sox5*, *Col6a2, Col12a1,* and *Chodl* in two comparable sections to those in Figure 4 from biological replicates. N=3 total.

**Supplementary File 1. Spatial feature plots for top 24 markers of cluster 4 in Figure 5 Visium analysis.**

**Supplementary File 2. Spreadsheet of all marker genes across clusters for WT1R1 section used for Figure 5 Visium analysis.**

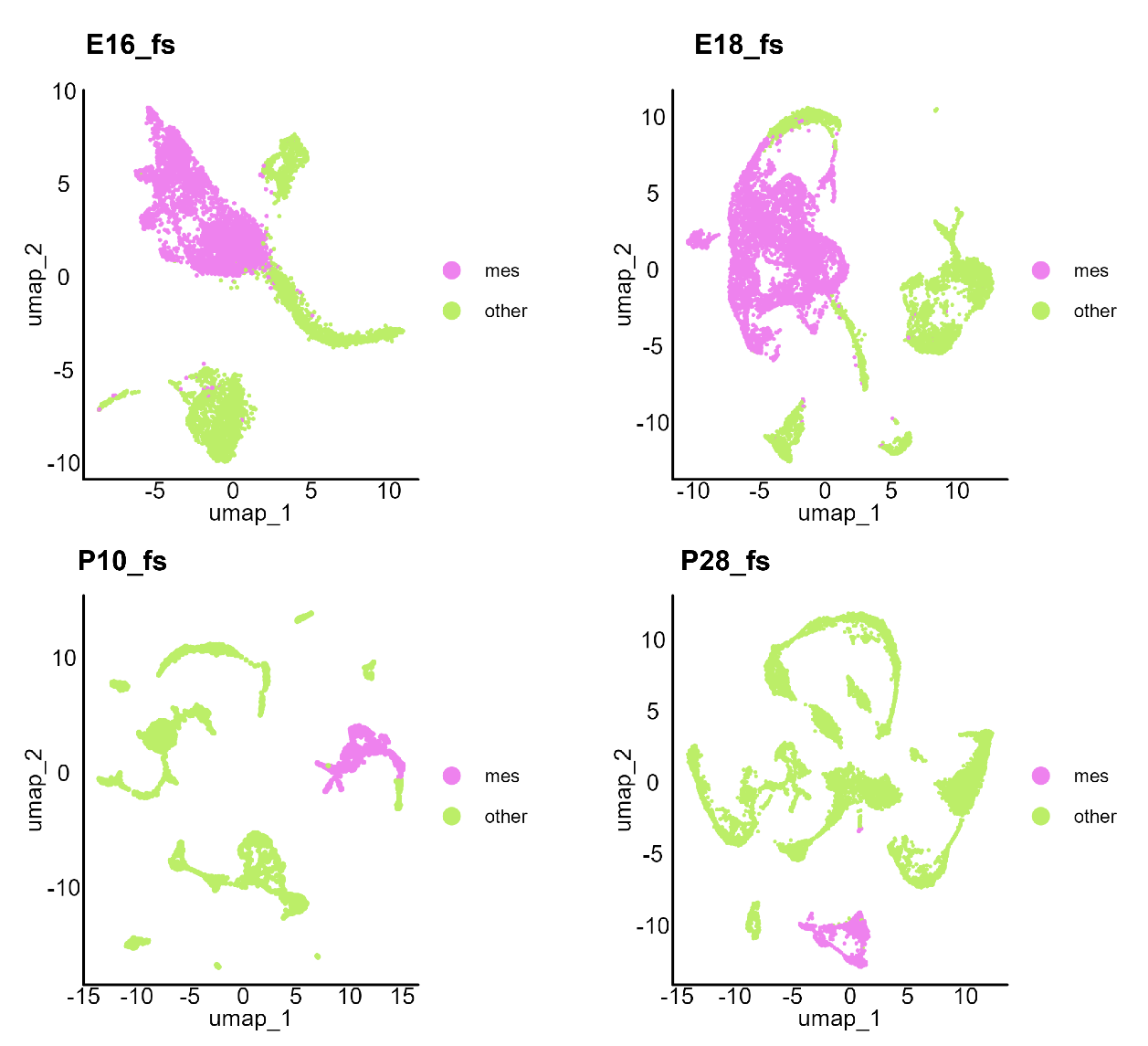

**Supplementary Figure 5. Grouped clustering of frontal suture single-cell RNA-sequencing UMAP into mesenchymal (mes) and non-mesenchymal (other) clusters as used in Figure 5 analysis.**

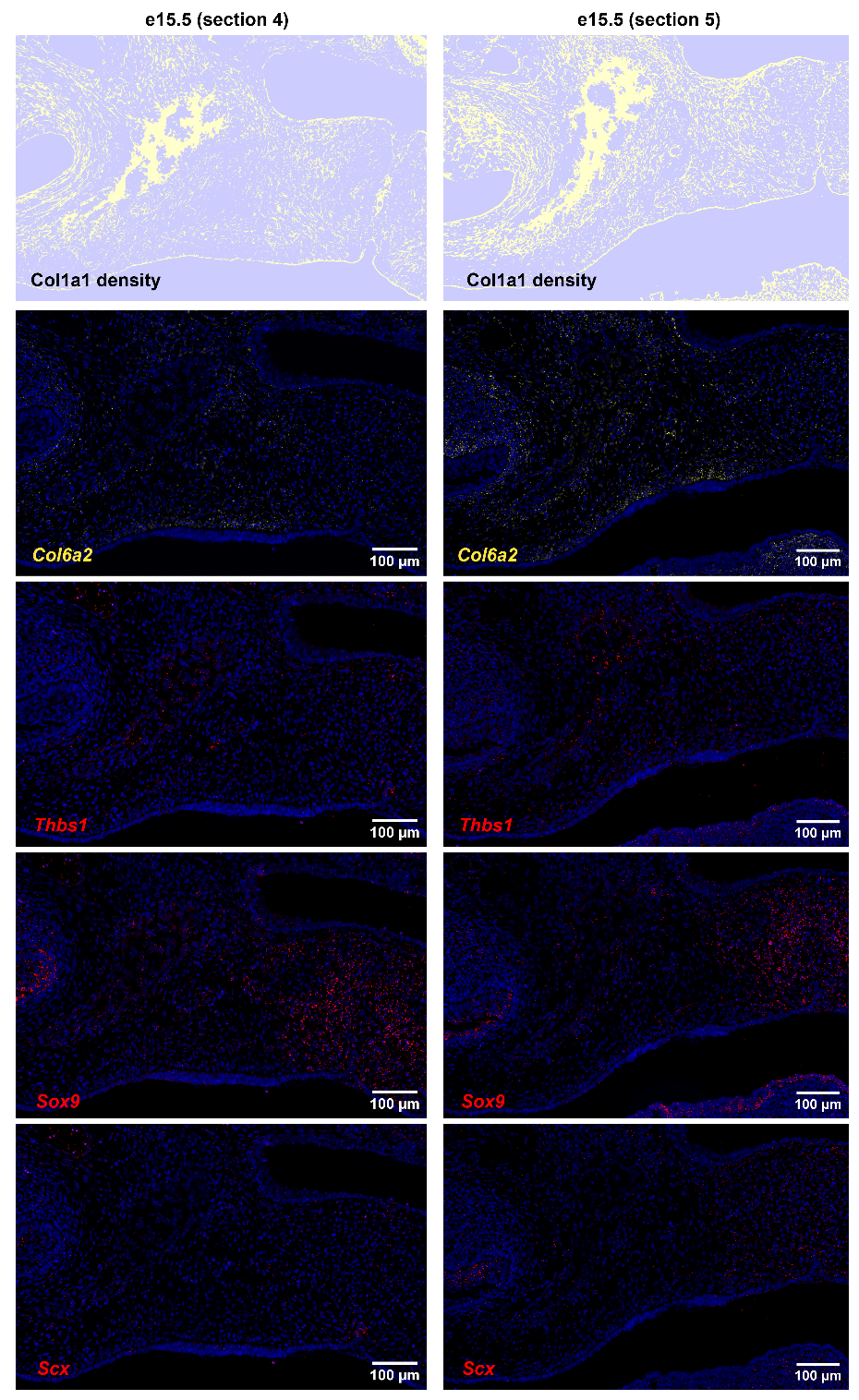

**Supplementary Figure 6.** *Col1a1 protein and* Col6a2, Thbs1, Sox9, Scx *mRNA detection with sequential RNAscope Hiplex hybridization and immunofluorescent staining.* Images in each column are from different mice, for a total sample size of 3 embryonic day 15.5 (e15.5) mice, including images shown in Figure 6.

**Supplementary File 3.** *Density of gene expression in the embryonic day 15.5 (e15.5) palatal shelf.* Collection of density gene expression plots for selected targets related to Bone Morphogenetic Protein, Fibroblast Growth Factor, Hedgehog, Matrix Metalloproteinase, Insulin-related Growth Factor, Transforming Growth Factor, and Wnt signaling networks. All images are from the Xenium experiment depicted in the main figure.
