## Supplementary figures and images for "Tendon-associated gene expression precedes osteogenesis in mid-palatal suture establishment"

### Supplementary File 1

#### FeaturePlots of top WT1R1 Visium Cluster 4 markers (e15.5 coronal)

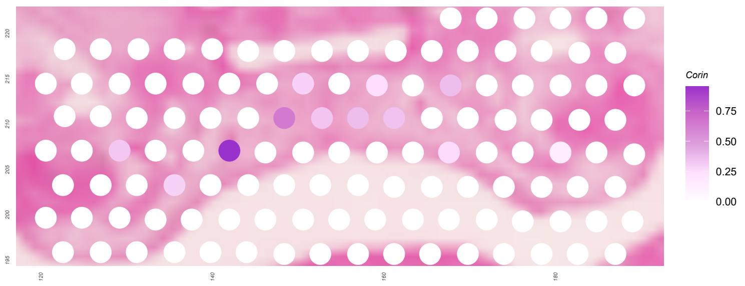


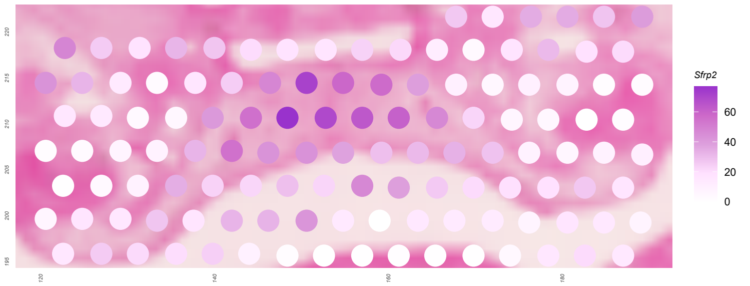


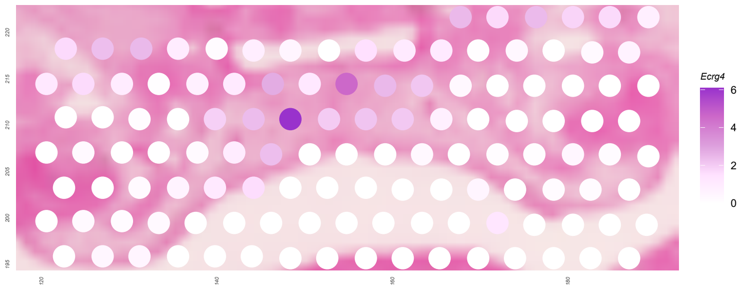


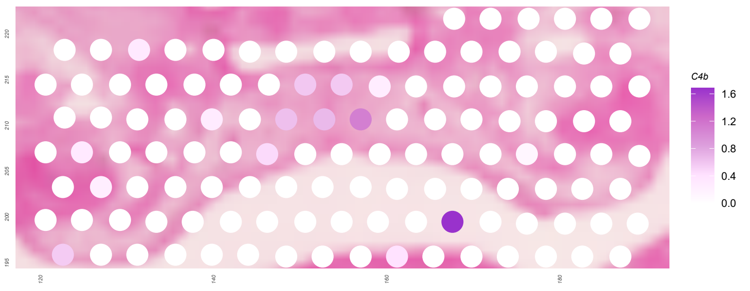


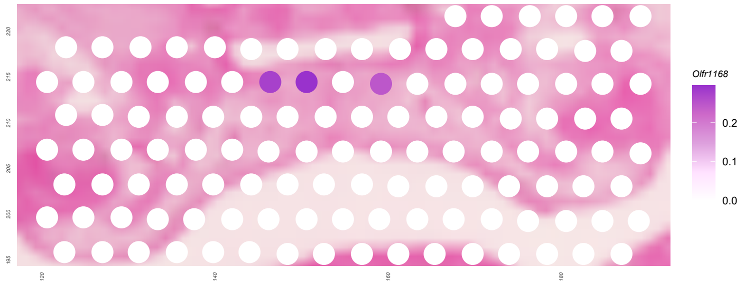


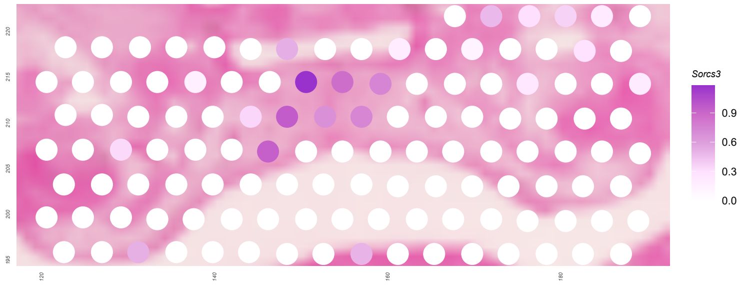


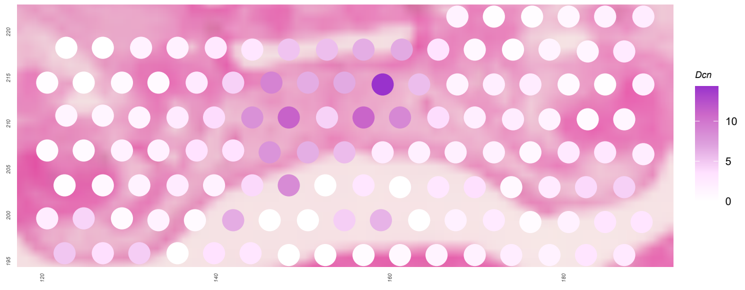


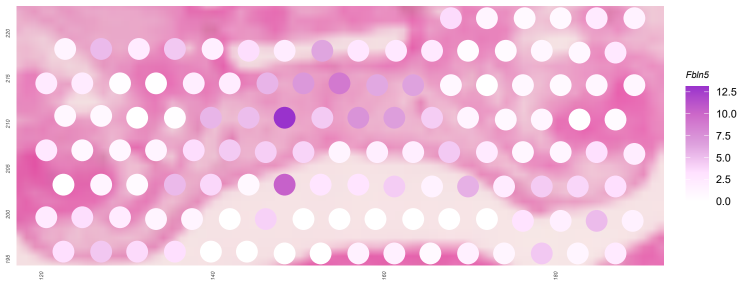

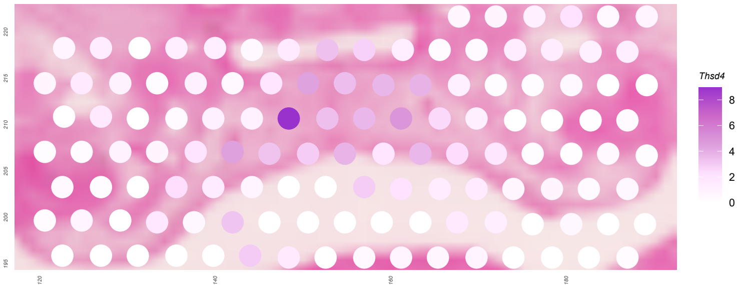

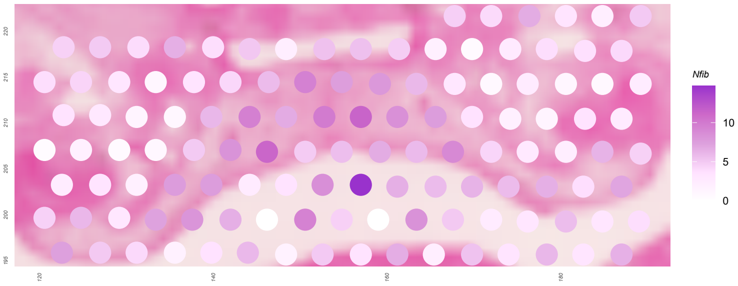

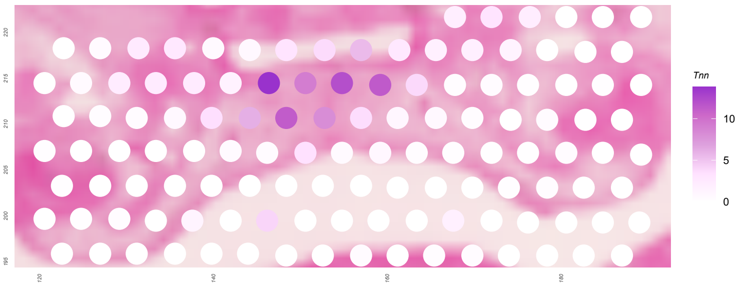

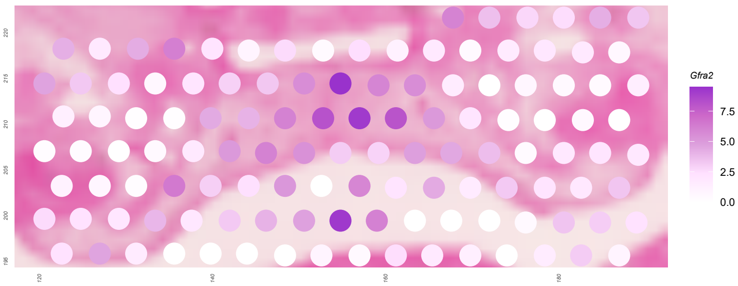

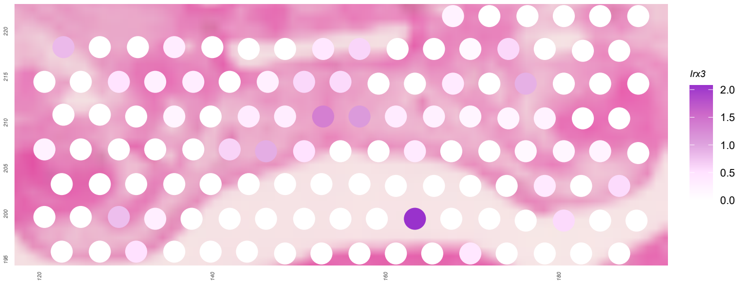

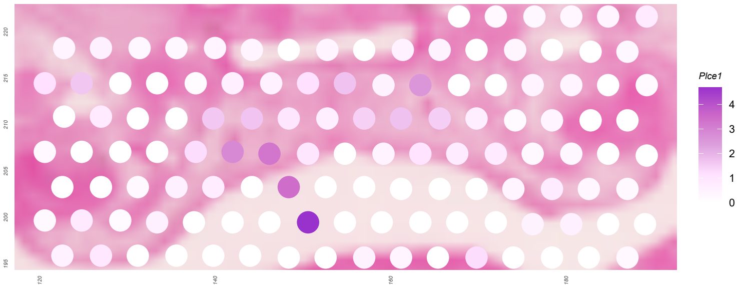

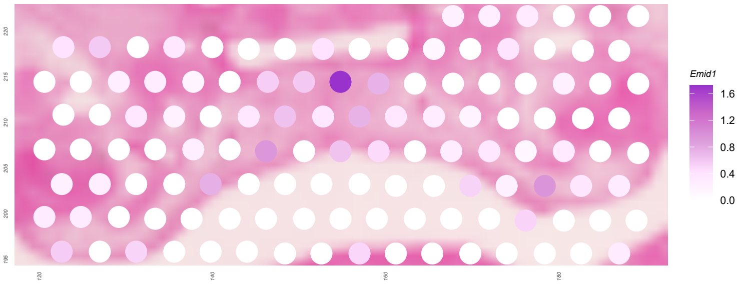

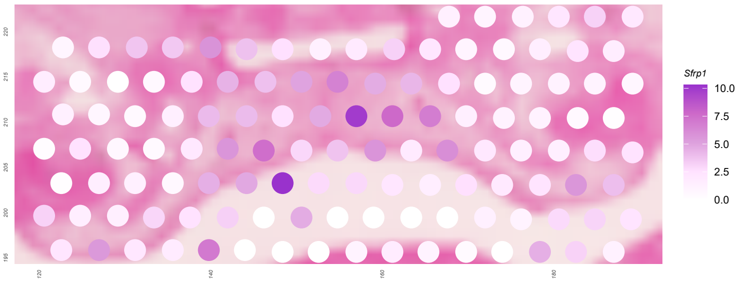

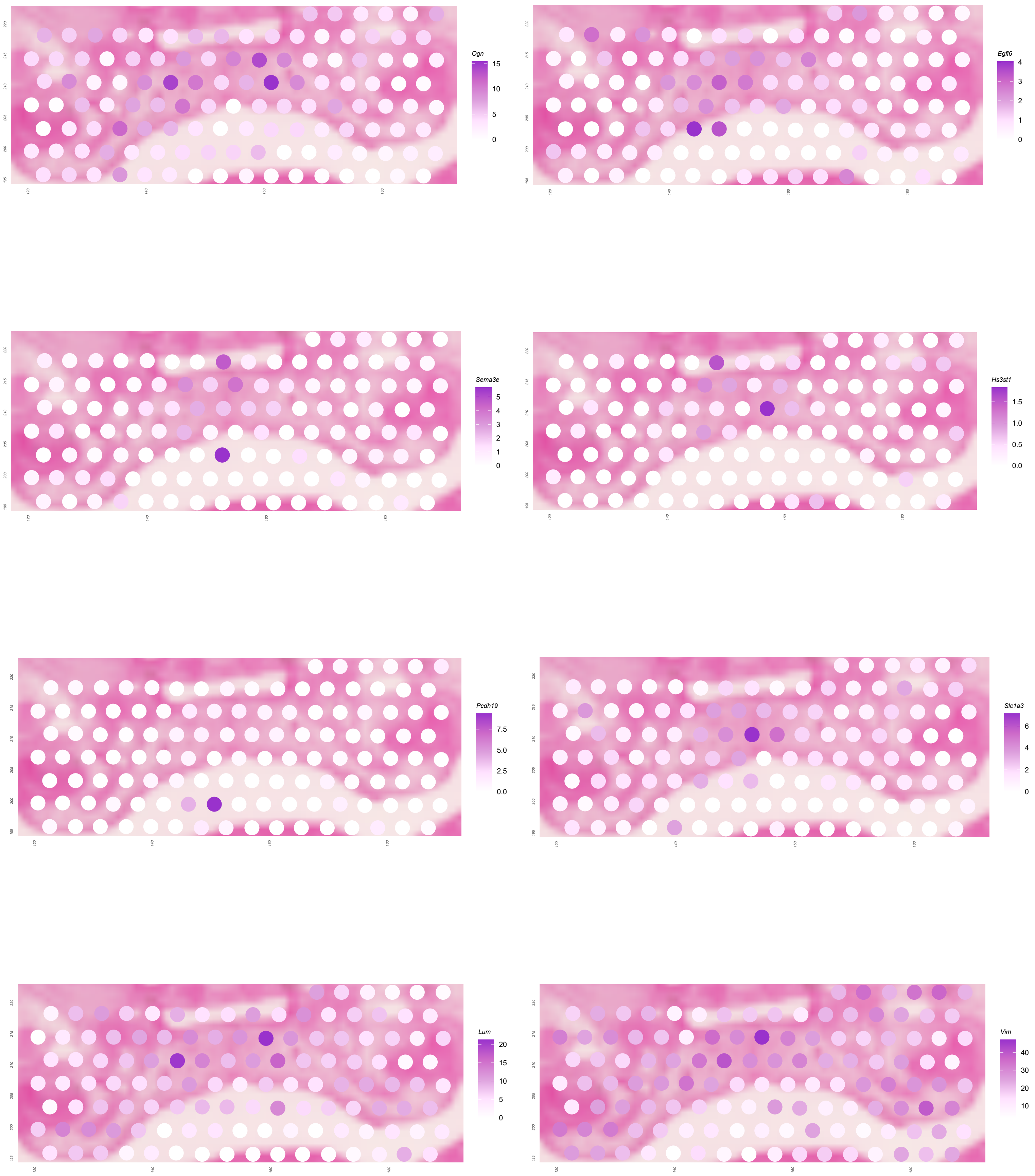
