## Supplementary File 3 for "Tendon-associated gene expression precedes osteogenesis in mid-palatal suture establishment"

**Bone Morphogenetic Protein signaling**

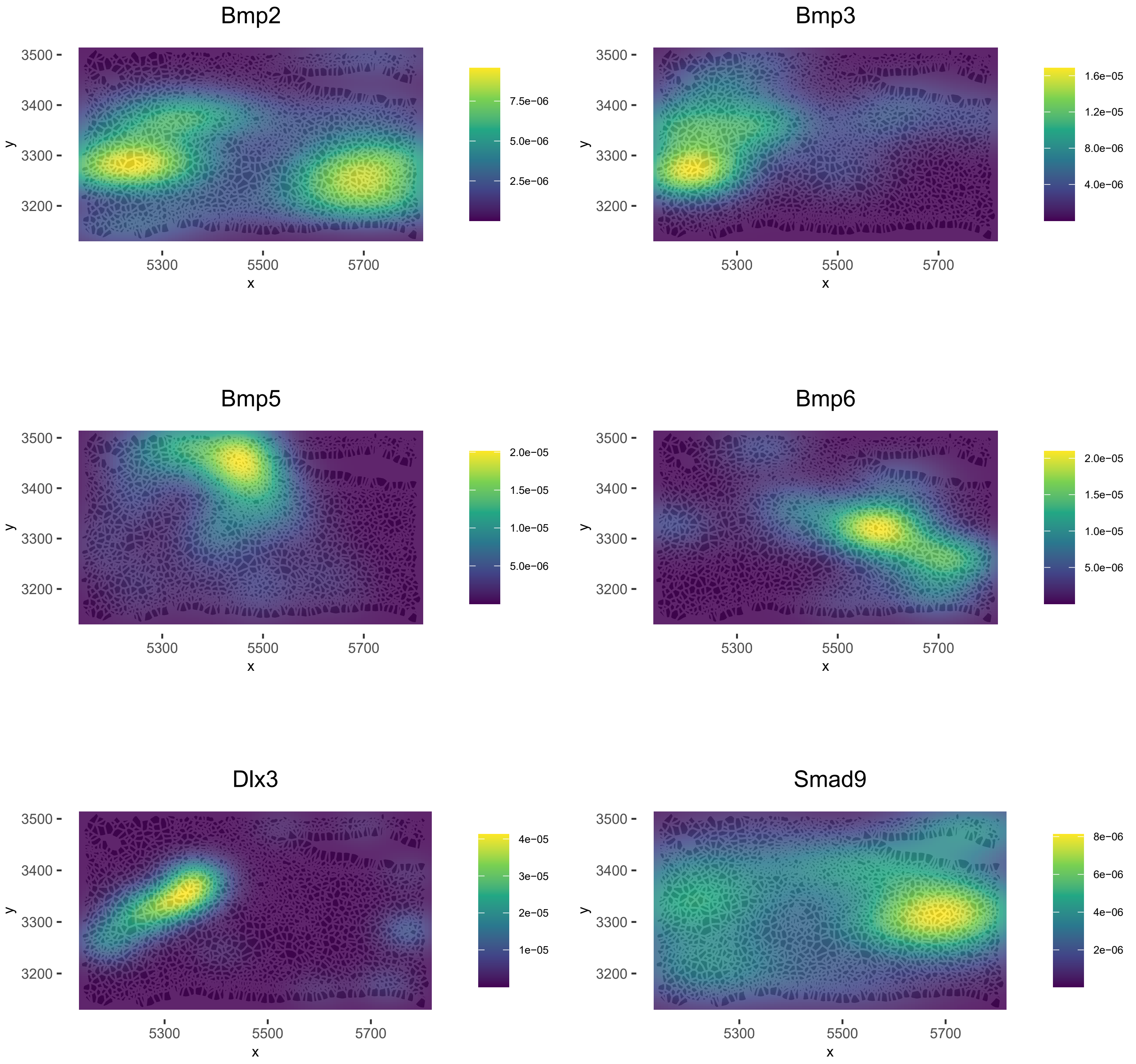

**Fibroblast Growth Factor signaling**

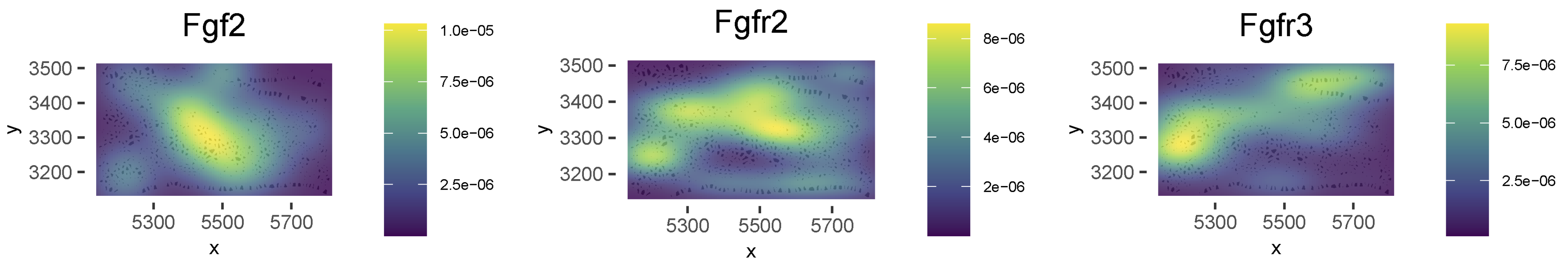


**Insulin-like Growth Factor signaling**

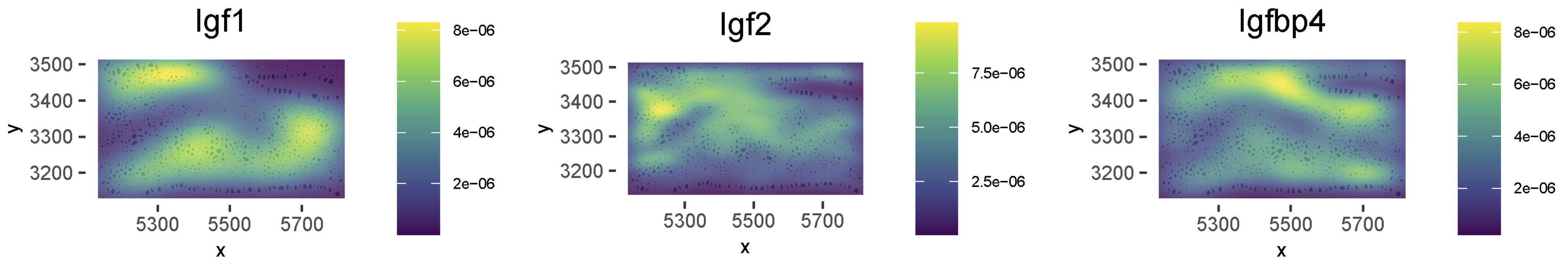

**Hedgehog signaling**

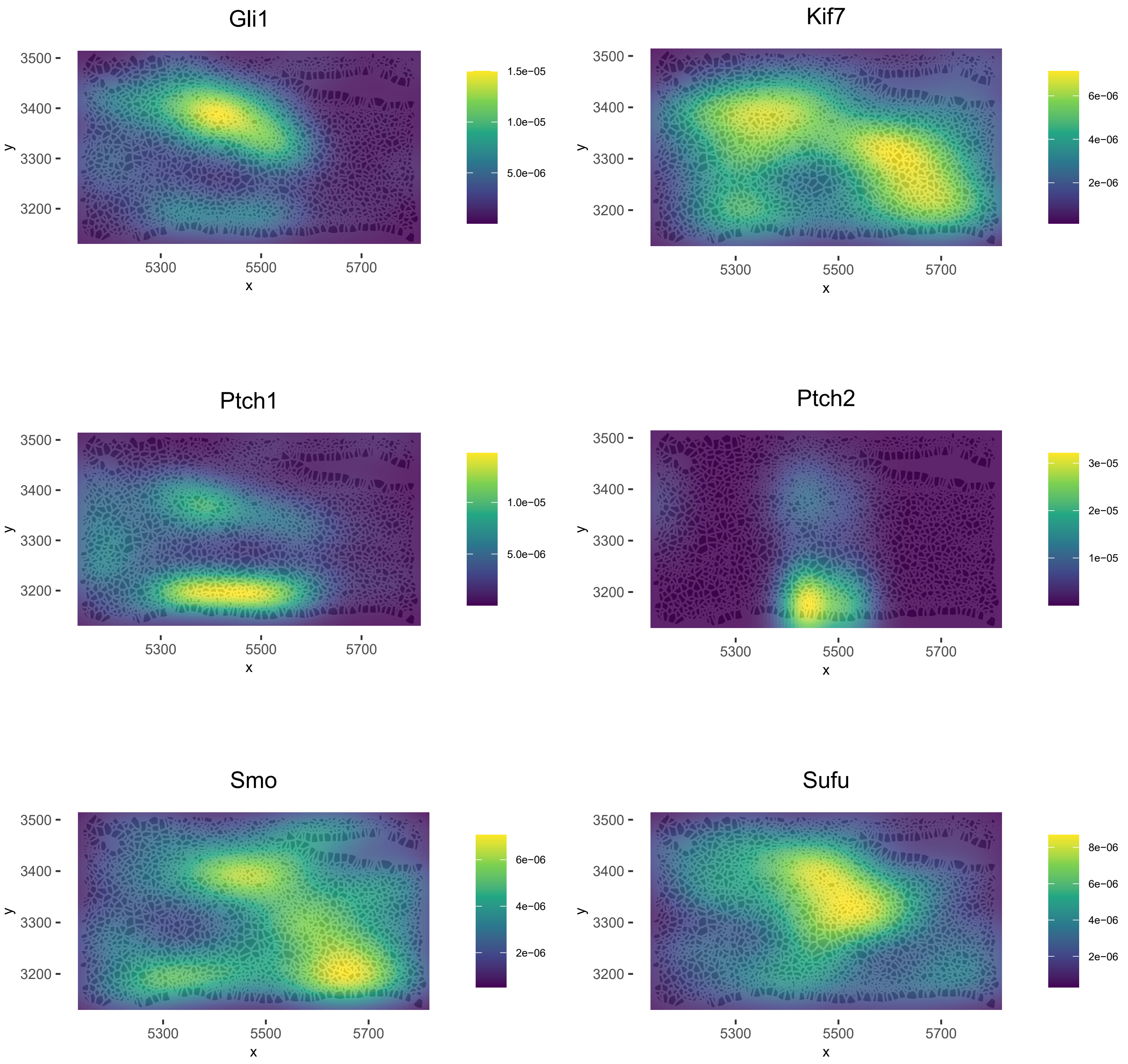

**Matrix Metalloproteinases**

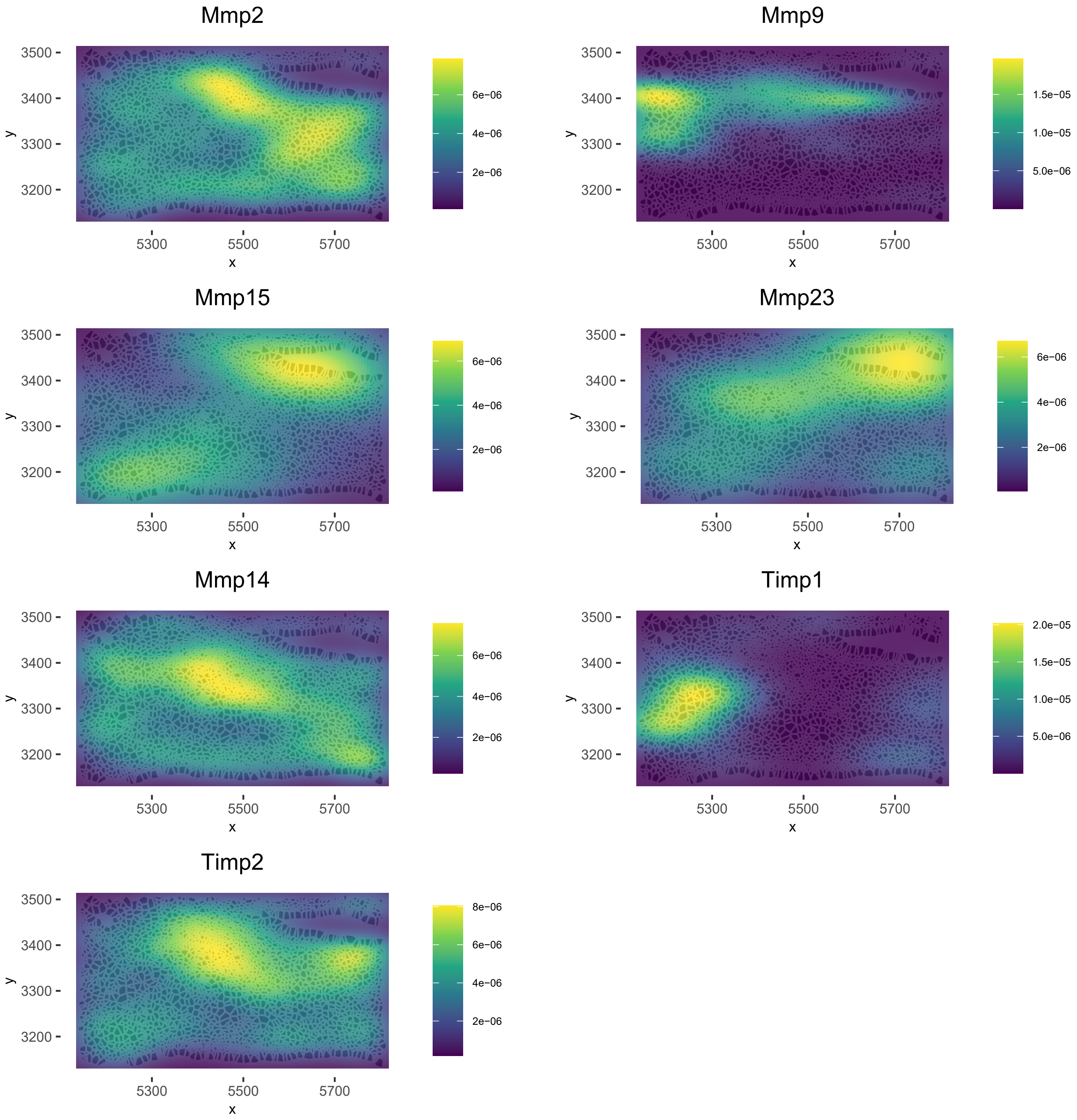

**TGF-beta signaling**

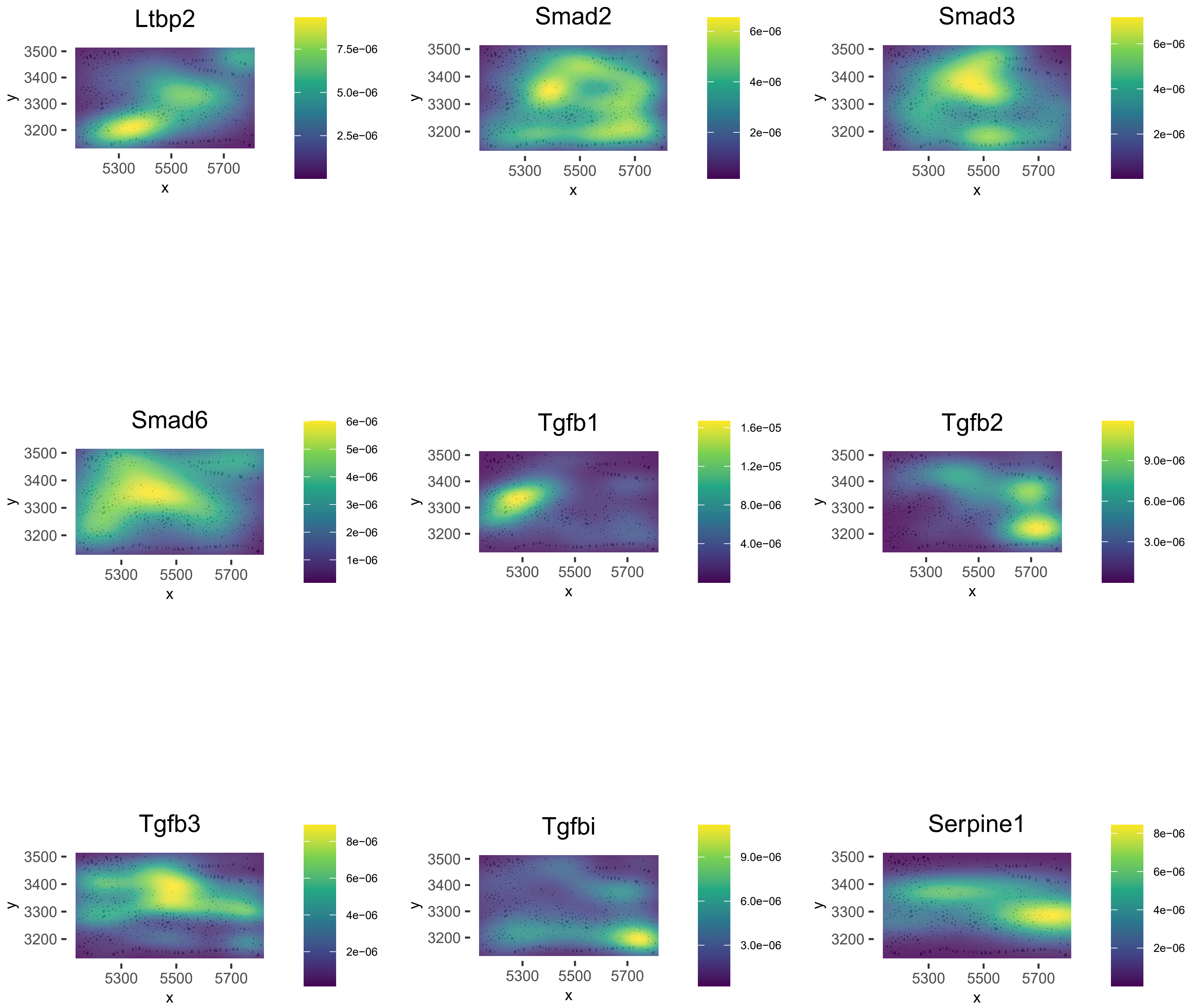

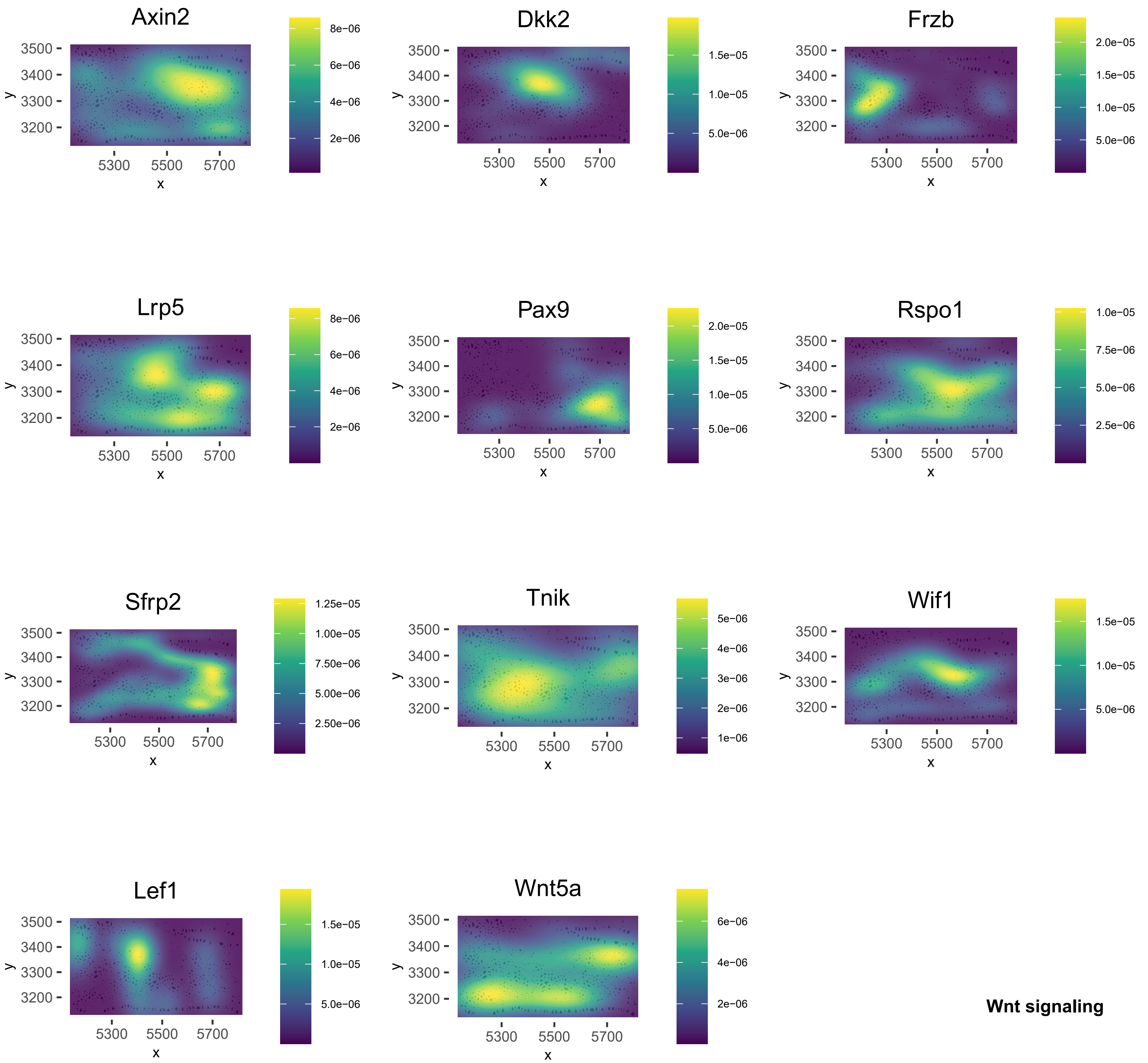
Axin2
Dkk2
Frzb
Lrp5
Pax9
Rspo1
Sfrp2
Tnik
Wif1
Lef1
Wnt5a
3500
3400
3300
3200
y
5300
5500
5700
x
8e−06
6e−06
4e−06
2e−06
1.5e−05
1.0e−05
5.0e−06
2.0e−05
1.5e−05
1.0e−05
5.0e−06
8e−06
6e−06
4e−06
2e−06
2.0e−05
1.5e−05
1.0e−05
5.0e−06
1.0e−05
7.5e−06
5.0e−06
2.5e−06
1.25e−05
1.00e−05
7.50e−06
5.00e−06
2.50e−06
5e−06
4e−06
3e−06
2e−06
1e−06
1.5e−05
1.0e−05
5.0e−06
1.5e−05
1.0e−05
5.0e−06
6e−06
4e−06
2e−06
Wnt signaling
